# Tissue-Like 3D Standard and Protocols for Microscope Quality Management

**DOI:** 10.1101/2022.08.14.503777

**Authors:** Benjamin Abrams, Thomas Pengo, Tse-Luen Wee, Rebecca C. Deagle, Nelly Vuillemin, Linda M. Callahan, Megan A. Smith, Kristopher E. Kubow, Anne-Marie Girard, Joshua Z. Rappoport, Carol J. Bayles, Lisa A. Cameron, Richard Cole, Claire M. Brown

**Affiliations:** Life Sciences Microscopy Center, 150 Sinsheimer Labs, University of California, Santa Cruz, 1156 High Street, Santa Cruz, CA 95064, USA, RRID:SCR_021135; Informatics Institute, University of Minnesota Twin Cities, Cancer and Cardiovascular Research Building, 2231 6th St SE, Minneapolis, MN 55449, USA; Advanced BioImaging Facility (ABIF), McGill University, 3649 Prom, Sir William Osler, Bellini Building, Room 137, Montreal, QC H3G 0B1, Canada, RRID:SCR_017697; Department of Physiology, McGill University, Montreal, QC; *St. Giles Foundation Advanced Microscopy Center, Cold Spring Harbor Laboratory, One Bungtown Rd., Cold Spring Harbor, NY, 11724, USA, RRID:SCR_023023; Department of Neuroscience, Del Monte Institute for Neuroscience, Univ. Rochester Medical Center, Rochester, NY 14642, USA; Biology Department, James Madison University, Bioscience Building, 951 Carrier Drive, Harrisonburg, VA 22807, USA, RRID:SCR_021904; Center for Genome Research and Biocomputing, Oregon State University, 1500 SW Jefferson Way Corvallis, OR 97331, USA; Center for Advanced Microscopy and Nikon Imaging Center, Feinberg School of Medicine, Northwestern Medicine, Northwestern University, Chicago, IL, USA; *Boston College, 140 Commonwealth Avenue, Chestnut Hill, Massachusetts, USA; Institute of Biotechnology, Cornell University, Ithaca, NY, USA; Light Microscopy Core Facility, Duke University, 4215 French Family Science Center, 124 Science Drive, Durham, NC 27708, USA; New York State Dept of Health/Wadsworth Center, Advanced Light Microscopy & Image Analysis Core Facility, 150 New Scotland Ave, Albany, NY 12208, USA, RRID:SCR_021104

**Author notes:** Corresponding author: Claire M. Brown, PhD; Associate Professor, Physiology; Director, Advanced BioImaging Facility (ABIF), Department of Physiology, Bellini Building Room 137A. Equal contributions as first authors (Benjamin Abrams, Thomas Pengo, Tse-Luen Wee). Equal contributions as principal authors (Richard Cole, Claire M. Brown).

**Keywords:** standard, reproducibility, resolution, quality control, point spread function (PSF), Signal-to-Noise-Ratio (SNR), confocal microscopy, fluorescence microscopy, 3D imaging, quantitative imaging

## Abstract

This article outlines a global study conducted by the Association of Biomedical Resource Facilities (ABRF) Light Microscopy Research Group (LMRG). The results present a novel 3D tissue-like biologically relevant standard sample that is affordable and straightforward to prepare. Detailed sample preparation and instrument specific image acquisition protocols and image analysis methods are presented and made available to the community. The standard consists of sub-resolution and large well characterized relative intensity fluorescence microspheres embedded in a 120 µm thick 3D gel with a refractive index of 1.365. The standard allows the evaluation of several properties as a function of depth. These include: 1) microscope resolution with automated analysis of the point spread function (PSF), 2) automated signal-to-noise-ratio analysis, 3) calibration and correction of fluorescence intensity loss, and 4) quantitative relative intensity. Results demonstrate expected refractive index mismatch dependent losses in intensity and resolution with depth but the relative intensities of different objects at similar depths were maintained. This is a robust standard showing reproducible results across laboratories, microscope manufacturers and objective lens types (e.g. magnification, immersion medium). Thus, these tools will be valuable for the global community to benchmark fluorescence microscopes and will contribute to improved rigor and reproducibility.

## INTRODUCTION

There has been a growing awareness of the importance of standards, rigor and reproducibility in science (Collins & Tabak, 2014; LaBaer, et al., 2018). The issue was recently highlighted through many articles in a feature Nature Methods issue highlighting *“Reporting and reproducibility in microscopy”* (Boehm, et al., 2021; Hammer, et al., 2021; Montero Llopis, et al., 2021; Nelson, et al., 2021; Swedlow, et al., 2021). It has been shown that a significant amount of published scientific data cannot be reproduced, in some cases even by the laboratory that originally performed the work (Franca & Monserrat, 2018). In addition, microscope methods reporting is weak overall so there is often no possible way that others can reproduce experimental results (Marques, et al., 2020). High-resolution confocal laser scanning microscopy (CLSM) is broadly used in many fields of science (Jonkman, et al., 2014; Jonkman, et al., 2020). However, routine quality management and assessment of instrument performance is rarely done. Quality management and assessment would provide a clear way to enhance rigor and reproducibility and should be a prerequisite for quantitative imaging-based research. The barriers to instrument performance testing include a lack of relevant, cost-effective test specimens, software to acquire and analyze the performance test data, time to dedicate to in-depth accurate calibrations and a standardized methodology for microscope users to collect, analyze and interpret the test results (Nelson, et al., 2021). Without rigor and reproducibility, science cannot confidently build on previous results. Validating scientific outcomes within and across different laboratories is a shared responsibility of all stakeholders involved in the scientific endeavor (Boehm, et al., 2021).

The Association for Biological Resource Facilities (ABRF) is committed to developing and disseminating best practices and standards (Knudtson, et al., 2019; Mische, et al., 2020). More Specifically, the Light Microscopy Research Group (LMRG) is focused on standards and protocols for reproducible light microscopy data collection. To that end, the ABRF-LMRG has conducted several studies which have sought to raise awareness in the community, develop and implement standard samples and image acquisition and analysis procedures to assess the essential aspects of microscope quality for quantitative imaging (Brown, et al., 2015; Cole, et al., 2011; Cole, et al., 2013; Deagle, et al., 2017; Stack, et al., 2011).

The guiding design principles behind these studies have been to reach a broad international audience, use samples that are affordable and easy to obtain or prepare, and use protocols that are straight forward and applicable across microscope platforms from all manufacturers. One inexpensive resource available for testing microscope performance is the fluorescent microsphere or bead (Goodwin, 2007). In 2011, the ABRF-LMRG developed protocols for preparing samples and measuring resolution of standard CLSM microscopes (i.e. not super resolution) with 100 nm diameter fluorescent microspheres (Cole, et al., 2011). The ABRF-LMRG also published study results focused on developing samples and protocols for measuring CLSM laser stability, co-registration between color channels and uniformity of illumination across the microscope field-of-view (Stack, et al., 2011). In a separate publication, a protocol was developed to automatically calculate field uniformity and determine if it needed to be corrected (Brown, et al., 2015). Further work by the group involved the development of samples and protocols for measuring resolution at the coverslip-sample interface, measuring spectral detector accuracy with mirror slides and used a two-color double orange microsphere sample (i.e. microspheres with an orange ring and orange core) to evaluate the accuracy of spectral unmixing/separation algorithms (Cole, et al., 2013). Finally, protocols have been published for quality control and general microscope cleaning and maintenance (Deagle, et al., 2017).

Many other groups have contributed to this field with protocols and software for benchmarking (Halter, et al., 2014) and calibration (Kedziora, et al., 2011) of widefield microscopes, quality control assessment of the confocal laser scanning microscope (CLSM) (Zucker, 2006a; Zucker, 2006b; Zucker & Price, 2001), automated software to monitor CLSM performance and track it over time (Hng & Dormann, 2013), evaluation of camera performance (Murray, 2013), evaluation of 3D fluorescence microscope performance (Murray, et al., 2007) and automated measurements of noise within CLSM images (Ferrand, et al., 2019). Several excellent reviews are also available covering multiple aspects of quantitative fluorescence imaging including CLSM (Jonkman, et al., 2014; Jonkman, et al., 2020; Lambert & Waters, 2014; Murray, 2013; Waters, 2009; Waters & Wittmann, 2014). More recently, the international **QU**ality **A**ssessment and **REP**roducibility for Instrument and Images in **Li**ght **Mi**croscopy (QUAREP-LiMi) has been established(Boehm, et al., 2021). QUAREP-LiMi is focused on broad activities related to microscope quality management through many focused working groups (Nelson, et al., 2021). Members of the ABRF-LMRG are actively participating in QUAREP-LiMi and the work presented here compliments the initiatives of this international network.

Most microscopy standards and protocols have been developed for 2D applications and there are very few 3D standards. These 2D samples can evaluate instrument performance but they lack the ability to provide direct information regarding performance through the depth of a biological tissue sample that can be tens to hundreds of microns thick. Proprietary commercial laser patterned test slides including 3D patterns are available from Argolight and PSFCheck (Corbett, et al., 2018). These materials do not have a refractive index similar to biological samples so cannot be used for direct calibration or correction of artifacts in thick biological samples. However, the biological specimen itself does impact quantitative imaging and data interpretation (Reiche, et al., 2022) emphasizing the need for biologically relevant standards. **The goal of this study was to develop an affordable easy to prepare 3D fluorescent sample that could mimic thick biological samples and be used to benchmark CLSM performance.** Test samples include sub-resolution microspheres to measure aberrations, **S**ignal-to-**N**oise **R**atios (SNR) and resolution and fluorescent microspheres of different intensities to validate quantitative intensity comparisons by relative intensity as a function of depth. These samples, combined with clear straightforward image acquisition and analysis protocols, provide the necessary tools to improve reproducibility and provide a method for ‘on-instrument’ quality control and early detection of performance issues.

To assess the ease of sample preparation, image acquisition and validation of the protocols, sample kits were developed and sent to imaging core facilities around the globe. Each kit included protocols and materials to prepare two samples, one to measure resolution, aberrations and SNRs and one to measure relative intensity as a function of depth. Study participants were asked to follow detailed protocols to prepare samples at their own institutions, image the samples and then upload the data for analysis by the ABRF-LMRG. Detailed protocols for sample preparation and image acquisition on most major CLSM manufacturer platforms were provided and are available on GitHub (https://github.com/orgs/ABRFLMRG/). The ABRF-LMRG team evaluated the 3D data sets to measure how each of the following metrics changed as a function of imaging depth up to 100 µm in the sample: resolution, spherical aberrations, SNR, fluorescence intensity and relative fluorescence intensities. This manuscript provides an overview of the protocols for preparing and imaging the 3D samples and provides image analysis protocols and software tools to evaluate the data.

## MATERIALS AND METHODS

### Sample kits

All study participants received a sample kit which included: 1) 500 µL of CyGEL^TM^ mounting media (BioStatus Ltd., Cat# Cy10500), 2) three microscope slides (FisherBrand, Cat# 12-552-3), 3) three double-sided 9 x 0.12 mm round well electron microscopy grade double sided adhesive spacers (Electron Microscopy Sciences, Cat# 70327-8S), 4) four 18 x 18 mm square coverslips (Fisherbrand, Cat# 12541A, No. 1.5 coverslips, 0.15-0.19 mm thickness), and 5) two tubes with fluorescent microsphere samples.

Sample #1 contained a 5.0 µL sample with four types of 2.6-2.7 µm diameter microspheres. Two different intensities of green InSpeck^TM^ microspheres (505/515, ThermoFisher, Cat# I-7219, Lot# 1772680) and two different intensities of red InSpeck^TM^ microspheres (580/605, ThermoFisher, Cat# I-7224, Lot# 1859236). The microspheres were from ThermoFisher kits that include microspheres ranging from 0.1% to 100% brightness. For these studies, the 3% and 30% relative intensity microspheres were used. In practice, the microspheres are not exactly 3% and 30% relative intensity as there is some batch-to-batch variability. However, ThermoFisher provides specification sheets for each kit and lot number (available at https://github.com/orgs/ABRFLMRG/). The microspheres are subjected to quality control and the intensities of each lot are measured using flow cytometry. **For the microspheres used in this study the relative intensities for the bright/dim green microspheres were 35% and 3.7% for a ratio of 9.5 and the bright/dim red microspheres were 40% and 4.7% for a ratio of 8.5.** The purpose of this sample was to measure changes in fluorescence intensity and relative intensities up to 100 μm into the sample.

Sample #2 contained 2.5 μL of a 1:5000 dilution of both 100 nm diameter green microspheres (505/515, ThermoFisher, Cat# F-8803) and 1 µm orange microspheres (540/560, ThermoFisher, Cat# F-8820) for a total sample volume of 5 μL. The large orange microspheres were added to the sample to provide a marker to easily focus on the sample. However, the point spread function (PSF) resolution measurements were made with the green 100 nm diameter sub-resolution microspheres.

### Standard Slide Preparation

Standard slides were prepared by study participants. Both samples were prepared on ice as CyGEL^TM^ (RI: 1.365) is a thermo-reversible solution that is a liquid at low temperature and a gel at room temperature and above. The RI is stable once the product transitions from a liquid to a gel and the sample is sealed or kept under high humidity. Microspheres embedded in CyGEL^TM^ should be prepared fresh each day to avoid drying out, formation of air bubbles and potential RI changes. The microsphere aliquots from the kit were sonicated for 15-20 min in an ice bath to break up any microsphere aggregates. The microsphere solution was then combined with 100 µl of ice cold CyGEL^TM^ and mixed gently by pipetting the mixture up and down slowly to avoid the formation of bubbles. **Note: it is *essential* to ensure the mixture stays cold throughout the entire protocol to keep the CyGEL^TM^ from solidifying prematurely.** In general, no more than 5 µL of microsphere solution should be added as the CyGel^TM^ will become watered down and will not properly solidify. Microscope slides were pre-washed with 70% ethanol and the EM spacer was firmly fixed onto the center of the slide. The upper adhesive side of the EM spacer was exposed before placing the gel-bead mixture on the slide. A micropipette was used to dispense 10 µL of the microsphere-gel mixture gently into the spacer well on the microscope slide. This was done on the lab bench at room temperature to ensure the gel would solidify quickly and the microspheres would not settle to the bottom of the sample. The gel was covered quickly with a precleaned (70% ethanol and flamed) 18 x 18 mm #1.5 coverslip (0.15-0.19 mm thickness). Higher tolerance coverslips with precisely 0.17 mm thickness can also be used. Gentle pressure was applied to the center of the coverslip using a cotton swab to move any air bubbles out towards the edge of the coverslip to be released into the air. **<u>Important Note:</u> The slide was then placed inverted in a 37°C incubator for 5 minutes to ensure complete gel curing and minimize microsphere settling to the bottom of the sample.** The coverslip was then sealed with 2-3 coats of quick dry nail polish (e.g. Sally Hansen, Insta-Dri) in order to prevent the gel from drying out. Slides were left inverted (coverslip down) to cure for 1-2 hours and then checked on a fluorescence microscope for even microsphere distribution throughout the 100 µm thick gel. A detailed video of the microsphere sample preparation can be found at https://www.youtube.com/watch?v=KjVHZaLE14E.

***NOTE: Once the sample slides are created, the samples must be kept at room temperature*** *If they are stored in the cold, the CyGel^TM^ will liquify and the microspheres will settle to the bottom of the sample. Prepared samples cannot be shipped as the temperature is too low during air transit, causing the microspheres to settle. Samples can be kept for 1-2 weeks at room temperature. After that, the gel begins to dry out and air bubbles form under the coverslip*.

### Image Acquisition

Participants collected data on CLSMs using instrument-specific protocols provided for many different confocal models covering the four major manufacturers, Leica, Nikon, Olympus and Zeiss (detailed protocols are available on the ABRF-LMRG GitHub page at https://github.com/orgs/ABRFLMRG/). Participants used a wide range of objective lenses and immersion media. Objective lens magnification ranged from 20x to 63x. Objective numerical aperture (NA) values ranged from 0.7 - 1.402 and immersion types included air, water, glycerol and oil.

For the resolution/PSF standard samples, high resolution settings were used to achieve a pixel size of ∼70 nm and z-spacing of 200 nm to enable accurate curve fitting and determination of the PSF size and shape in 3D. For lower resolution objective lenses larger sampling frequencies can be used. Recommended settings are available in Table 1 of Cole et. al. 2011. Images were collected at 8-bit to keep data set sizes small to allow for analysis of many microspheres in a large sample volume using the image analysis pipeline.

**TABLE 1:** Lateral (top) and Axial (bottom) FWHM as a function of depth into the 3D standard sample relative to theoretical objective resolution. Quantitative summary of the FWHM values from the PSF measurement data of 100 nm diameter sub-resolution green microspheres. Values were categorized by objective lens NA, immersion medium and depth quadrant, “Q: (see Fig. 2 legend for details). The “Difference in Median Value” column shows the difference in median lateral (top table) or median axial (bottom table) PSF value between depth quadrants 1 and 4. The Theoretical Resolution values were calculated based on the formula given and referenced in the materials and methods section. The Bulk Population Mean and Bulk Population Standard Deviation columns refer to the data presented in figures 3A and 3B, where the black horizontal line is drawn at the mean value.

| Summary of Lateral FWHM ($\mu\text{m}$) Measurements | | | | | | | |
| --- | --- | --- | --- | --- | --- | --- | --- |
| NA | Difference in Median Value Q1:Q4 | Difference in Median Value Q1:Q4 as a Percentage of Theoretical Resolution | Theoretical Resolution | Bulk Population Mean | Bulk Population StDev | Difference Between Bulk Population Mean and Theoretical Resolution | Difference Between Bulk Population Mean and Theoretical Resolution as a Percentage |
| 0.7 | 0.008 | 2% | 0.355 | 0.430 | 0.043 | 0.075 | 21% |
| 1.2 | -0.008 | -4% | 0.207 | 0.308 | 0.053 | 0.101 | 49% |
| 1.25 | 0.036 | 18% | 0.199 | 0.280 | 0.032 | 0.081 | 41% |
| 1.3 | 0.001 | 1% | 0.191 | 0.277 | 0.038 | 0.086 | 45% |
| 1.4 | 0.050 | 28% | 0.178 | 0.296 | 0.042 | 0.118 | 66% |
| Summary of Axial FWHM ($\mu\text{m}$) Measurements | | | | | | | |
| NA | Difference in Median Value Q1:Q4 | Difference in Median Value Q1:Q4 as a Percentage of Theoretical Resolution | Theoretical Resolution | Bulk Population Mean | Bulk Population StDev | Difference Between Bulk Population Mean and Theoretical Resolution | Difference Between Bulk Population Mean and Theoretical Resolution as a Percentage |
| 0.7 | 0.144 | 10% | 1.502 | 2.29 | 0.286 | 0.788 | 52% |
| 1.2 | -0.165 | -29% | 0.567 | 1.30 | 0.312 | 0.733 | 129% |
| 1.25 | 0.944 | 145% | 0.652 | 1.25 | 0.395 | 0.598 | 92% |
| 1.3 | 0.744 | 136% | 0.548 | 1.16 | 0.350 | 0.612 | 112% |
| 1.4 | 0.872 | 190% | 0.459 | 1.24 | 0.405 | 0.781 | 170% |

The intensity ratio standard samples were imaged with the CLSM setup for imaging a green dye (similar to EGFP, Alexa488 or FITC) and/or a red dye (similar to mCherry, Alexa594, or TRITC), see instrument specific protocols for details (https://github.com/orgs/ABRFLMRG/). Images were collected in 12-bit format for maximum precision in intensity analysis. For 2.6-2.7 µm diameter microsphere imaging, the pixel size in the software was set to 200-300 nm in x, y with a z-spacing of 1.0 µm.

For both standard samples, line averaging of 4 was used to balance a good SNR and time to collect 3D datasets, the photomultiplier (PMT) gain was set to achieve strong signal without saturating, the digital offset/background was set to ensure all pixels had a positive intensity value and that no pixels measured zero intensity to avoid cutting off low intensity data. All DIC related components were removed from the light path, and the pinhole was set to 1 Airy Unit (AU). Images submitted to the ABRF-LMRG were verified before analysis to ensure no pixel intensity values were saturated.

### Image Analysis

#### Part 1: Microscope Resolution Analysis with Sub-resolution Microsphere Samples

Data from Sample #2 preparations were submitted by study participants as 3D z-stacks of 100 µm depth of 100 nm green microspheres in CyGel^TM^. Data were collected with objective lenses of different magnification, numerical aperture (NA) and immersion media combinations. Microscope resolution as a function of depth into the CyGel^TM^ was measured via changes in the size of the Point Spread Function (PSF) of many 100 nm diameter green, fluorescent microspheres through the 100 µm sample depth. The initial 3D PSF analysis was performed with a modified version of the ImageJ plugin *PSFj* (Theer, et al., 2014). Each data set included hundreds of microspheres and more than one hundred datasets were received for analysis. Thus, the *PSFj* code was modified to manage batch processing of the high number of 3D volume image data sets. The *PSFj* code was later adapted and re-implemented with new code to enable analysis of multiple datasets in MATLAB (MATLAB 2017b, Mathworks, Natick, MA). The `findpeaksscript` MATLAB code (available at https://github.com/orgs/ABRFLMRG/) was designed to run the PSF calculations in parallel across multiple datasets, retained the user-reported metadata with the images throughout the analysis pipeline and automated PSF curve fitting data output in a format that could be easily analyzed, combined, and processed. Due to the high number of large 3D image datasets, the analysis was performed on a large-memory node hosted at the University of Minnesota Supercomputing Institute (dual 14-core E5-2650 Xeon processor, 1TB memory). Individual facilities should be able to run the analysis code in their facilities on a standard workstation equipped with MATLAB.

Briefly, the MATLAB algorithm finds local intensity maxima, estimates the z-position of the focal plane for each peak (see intensity peak analysis section below) and fits the raw unsmooth intensity of the x, y image of the microsphere in that focal plane to a 2D Gaussian *G*(*x*, *y*) with seven free parameters *I_b_* =background intensity, *I_0_* =peak intensity amplitude *x_0_*, *y_0_* = peak position 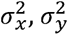 = intensity standard deviation in *x* and *y*, *θ* = rotation).

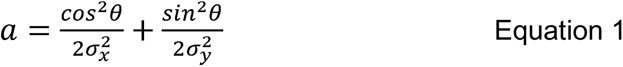

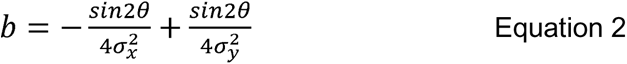

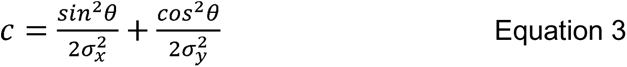

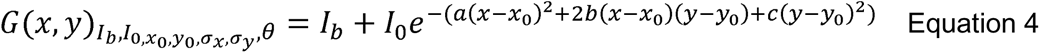

MATLAB algorithm: Levenberg-Marquardt (LM) algorithm, as implemented in the MATLAB function “*lsqcurvefit*” (https://www.mathworks.com/help/optim/ug/lsqcurvefit.html).

##### Intensity peak Identification

Each image was first filtered using a separable binomial filter with kernel [1,8,28,56,70,56,28,1]/248 in each dimension (roughly equivalent to a 2D Gaussian filter *σ* = 1 pixel or 70 nm). The filtering procedure reduced noise in the image without affecting the estimate of the microsphere location. A 3D regional maxima operation was applied to the smoothed volume to highlight microspheres identified as a local intensity higher than the surrounding area. A 3D connected components analysis then labeled each microsphere and calculated the centroid in x, y and z and the mean intensity. Only spots whose average intensity was 3 times higher than the mode of the image (i.e. the background) were analyzed further.

##### Intensity peak analysis

Each intensity peak was analyzed individually. First, the intensity peak’s focal plane in z was estimated by fitting a one-dimensional Gaussian function to the vector of raw intensities at the centroid’s position in *x* and *y*, across a range of z-stack images above and below the maximum. The full width at half maximum (FWHM) in z was also calculated at this point with the formula 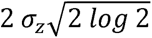 where *σ_z_* is the *σ* of the fitted Gaussian. A five-parameter Gaussian was fit to the array of raw intensities around the peak intensity x- and y-positions, at the z focal plane. Pearson’s correlation coefficient (R^2^) was calculated between the data and the curve fit and saved along with the fitted parameters for quality control purposes. The FWHM along the x, y and z-axes, the asymmetry (ratio of the z-axis FWHM over the x, y-axis FWHM), and the residual norm were saved. All curve fitting was performed using the Levenberg-Marquardt (LM) algorithm, as implemented in MATLAB function `lsqcurvefit`. Fit peaks that were less than the FWHM away from the intensity peak in both x and y and had an R^2^ above 0.9 were defined as valid, consistent with *PSFj*’s original criteria (Theer, et al., 2014).

##### PSF data filtering

To ensure the image acquisition protocol was followed properly (no zero intensity pixels) data filtering included selecting microspheres with local backgrounds between -10 and 10 corresponding to ∼4% of the maximum intensity in the 8-bit images.

Amplitude values with a range up to 4095 were allowed for the inclusion of datasets with 12-bit images, following the rationale that 12-bit data acquisition had not been made mandatory solely for data handling purposes and higher intensity pixels within a dataset would not affect the PSF shape. Peaks that were shifted by more than 1 pixel in either x or y from the initial regional maxima estimate were eliminated and only depth values between 0-100 μm were kept. FWHM has been a long respected standard, y and z-axis so FWHM values were also used for filtering. Data was retained if lateral FWHM was between 0.1-0.88 μm and axial FWHM values between 0.1-3.52 μm. These criteria were chosen to ensure that the fit metrics describing the PSF shape matched the data accurately. Three datasets per objective lens NA category was imposed to ensure sufficient data. Datasets were only included if they had at least 100 valid PSF measurements representing a minimal average microsphere density of one microsphere per micron of imaging depth.

In total, 140 datasets and 671,153 microsphere intensity peaks were analyzed. The number of datasets was reduced to 84 with a total of 17,781 peaks after removing images that set the offset/black level incorrectly leading to many pixels reading zero intensity units. The depth filter further reduced the number of data sets to 83, and the total number of peaks analyzed to 17,634. Amplitude filtering brought the number of peaks down to 17,552 over 68 datasets. Peak to fit displacement filtering lowered the total number of peaks analyzed to 17,547 over 68 datasets. FWHM filtering reduced the number of peaks to 17,494 over 68 datasets. Axial FWHM filtering brought the number of peaks down to 17,121 in 67 datasets. Requiring that each dataset have 100 peaks brought the number of datasets down to 35 containing a total of 15,912 peaks. The final filter requiring 3 datasets per NA category brought the total number of datasets analyzed to 30 with a combined total of 13,943 peaks, this data is presented in the following results section.

The theoretical lateral and axial resolutions were calculated using the following formulas as used in Cole 2011 (Cole, et al., 2011), originally from Carl Zeiss (Wilhelm, et al., 1997):

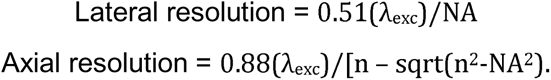

Where, λ_exc_ = wavelength of excitation n = refractive index of the immersion medium, NA = numerical aperture of the objective lens.

##### Signal-to-Noise-Ratio (SNR) Measurements

The SNR was calculated from image datasets from Sample #2 of the 100 nm sub-resolution microspheres. The noise was calculated for the intensity data points above the cutoff for the FWHM for each intensity peak as compared to the intensity curve fit values. The SNR was calculated by dividing the amplitude of the intensity peak by the standard deviation of the difference between the intensity data and the fitted Gaussian curve, restricting the calculation to the top half of each fitted peak (all x-y positions above the FWHM of the fitted Gaussian).

#### Part 2: Large Green and Red Microsphere Intensity and Relative Intensity as a Function of Depth

##### Initial Large Green and Red Microsphere Data Pruning

Image files from participants were opened with the Imaris 3D/4D Visualization and Analysis Software (Bitplane, Version 8.2.1, Oxford Instruments). Open-source software such as Fiji/ImageJ could be used to perform a similar analysis. The image properties, such as voxel dimensions and imaging depth (z-stack size) provided by the study participants and/or within the metadata for each dataset, were cross-referenced with the specifications outlined by the image acquisition protocols. Files were visually screened using the 3D viewing tool to verify that green (505/515) and red (580/605) microspheres of both low and high intensity were present throughout the image stack and that there was no data clipping (i.e. pixels reading zero intensity or saturated intensity). Datasets that did not meet these criteria were excluded from analysis.

##### Background Intensity Correction

The *Surface* function in Imaris was used to create a 3D **R**egion **O**f **I**nterest (ROI) around a green or red microsphere. The surface threshold was set using the automated k-means clustering based Imaris algorithm that identifies two classes of voxels with or without intensity signal. The surface of one microsphere was identified with a 3D ROI box and then the box was moved to a position within the dataset that did not include any surfaces (i.e. no intensity signal, no microspheres). In the statistics tab, the mean intensity of the background within the ROI for the green and red channels was measured. The mean background intensity was then subtracted from all voxel values in the green or red image stacks, respectively. These image stacks were saved as background intensity corrected image stacks and used for further quantitative intensity analysis as follows.

##### Fluorescence Intensity Data

The *Spot* function in Imaris was used to measure the fluorescence intensity data for all microspheres in each background intensity corrected image stack. The *Spot* function allowed specific and consistent x, y and z-diameters to be applied to each spot in the image file. This ensured consistency in intensity data measurements both within, and between, datasets. The datasheet for the specific InSpeck^TM^ microspheres used indicated the average microsphere diameter was 2.6 µm for the red microspheres and 2.7 µm for the green microspheres (see GitHub https://github.com/orgs/ABRFLMRG/ for detailed specification sheets). A value of 2.7 µm was used as the diameter in x, y for the Imaris *Spot*s analysis. The z-value for the *Spot*s was calculated to be twice this value (5.4 µm) to accommodate for the PSF signal elongation that is characteristic of 3D imaging. Using a value three times larger did not add significant intensity to the microsphere analysis but increased the chances of picking up additional nonspecific intensity from a nearby microsphere. So a z-value of twice the x, y-value maximized the intensity measured and minimized artifacts from nearby microspheres.

To begin *Spot* construction, the *Surpass* and *3D View* icons were selected and then the *Spot*s icon was chosen to open the *Creation Wizard*. Here the Imaris automatic algorithms were implemented and customized as needed. Automatic Imaris default selections in the *Creation Wizard* were left unchanged unless otherwise specified. In Step 2, the *Source Channel* was set to the green channel for the green (505/515) microspheres. The *Estimated x, y Diameter* was set to 2.7 µm, and the *Model PSF-elongation along z-axis* option was selected and 5.4 µm was input as the estimated *z-Diameter*. Additionally, *Background Subtraction* was unchecked, as the images were already corrected for background intensity. In Step 3, any automatic filters applied by Imaris were deleted. A filter was added to remove all the microspheres that were too close to the border/edge of the 3D volume to accurately measure the mean intensity. Due to the intensity differences between the dim (3.7%) and bright (35%) intensity microspheres, only the higher intensity microspheres were automatically detected by Imaris. The lower boundary of the automatic threshold set by the Imaris k-means clustering algorithm was then manually adjusted to also include the dim microspheres in the spot analysis dataset. If necessary, the *Display Adjustment* window was utilized to modify the image contrast to verify that all low intensity microspheres were identified as spots. Once the threshold was set appropriately, the *Spot*s construction was complete. The *Creation Parameters* were saved as ‘Spots 1X’ for future use and the completed *Spot*s for the green microspheres were saved as ‘Spots 1X – Green.’ Any microspheres that were clustered or overlapped with other nearby microspheres were excluded from the results using the manual *Delete* function in the *Edit* window.

The protocol outlined above was repeated for the red (580/605) dim (4.7%) and bright (40%) microspheres using the saved analysis parameters (Spots 1X). The saved parameters were loaded in Step 1, under *Favorite Creation Parameters.* The only modification for the red microspheres was in Step 2, where the *Source Channel* was set to the red channel. Completed *Spot*s for the red microspheres were saved as ‘Spots 1X – Red.’ The lower intensity threshold was again adjusted manually to include the dim microspheres as spots for the analysis.

The following variables for data export were selected for both the green and red microsphere intensity data: *Position x, Position y, Position z, Intensity Mean,* and *Intensity StdDev.* The data for the microspheres was exported from ‘Spots 1x – Green’ and ‘Spots 1x – Red’ from their respective raw image intensity channels. These data were sorted by their z-position and separated into depth quadrants of 25 µm increments (Q1: 0-24.99 µm, Q2: 25 µm-49.99 µm, Q3: 50 µm-74.99 µm, Q4: 75 µm-100 µm). The data was exported to a Microsoft Excel worksheet and sorted by *Intensity Mean* to separate the dim and bright intensity microspheres. To calculate the bright/dim intensity ratios and compare between datasets, microsphere intensities were normalized to the brightest microsphere in Q1 for each green or red data set.

Data visualization and plot creation was done using JMP Software by SAS, for all figures, except Figure 5. Figure 5a was created in Imaris by Bitplane and 5b was created in FIJI/ImageJ.

## RESULTS

### Test Kit Development, Distribution and Data Pruning

As with previous ABRF-LMRG studies, this study was announced on the confocal listserv (http://lists.umn.edu/cgi-bin/wa?A0=confocalmicroscopy) and at several microscopy focused meetings. Significant time was spent on trial-and-error to develop the ideal 3D ‘tissue mimic’ standard sample that would be affordable, robust and reproducible. The group settled on the use of a unique product, CyGel^TM^ from a study corporate partner, BioStatus Ltd. The product is in liquid form at low temperature and gel form at room temperature. It was developed to immobilize small organisms, cells or spheroids for long term live cell imaging but was also ideal for development of the 3D microscopy standard reported here. To minimize sample-to-sample variation, the original idea was to prepare all the samples with a small team and then ship them out to study participants. However, during shipping, the samples cooled, the CyGel^TM^ liquified and the microspheres settled to the microscope slide or coverslip surface and were no longer well distributed throughout the 3D sample. In the end, a second round of samples were sent out as test kits which contained all of the individual components and instructions on how to prepare samples. Detailed instructions and a link to a sample preparation video can be found at: https://github.com/orgs/ABRFLMRG/. Fluorescent microspheres were provided at no charge by ThermoFisher/Molecular Probes or The University of Minnesota shared multi-scale microscopy facility. CyGel^TM^ was provided at no charge by BioStatus Ltd. Test kits were sent out to Canadian and USA participants at no charge thanks to financial support from the ABRF. For international participants, BioStatus Ltd generously provided logistical and financial support and sent out test kits.

Test kits were sent out to 68 laboratories. A total of 191 image datasets were collected from 57 laboratories across 15 countries (Fig. 1A). Datasets from Confocal Laser Scanning Microscopes (CLSMs) from many major microscope manufacturers were received for analysis (Fig. 1B). Not all datasets were used for detailed analysis as they went through an initial pruning to eliminate poor quality data (see details in the Materials and Methods section). For example, image data sets were removed if they showed intensity saturation, low signal or if there were not enough microspheres imaged throughout the 100 µm depth of the 3D sample.

**Figure 1.**
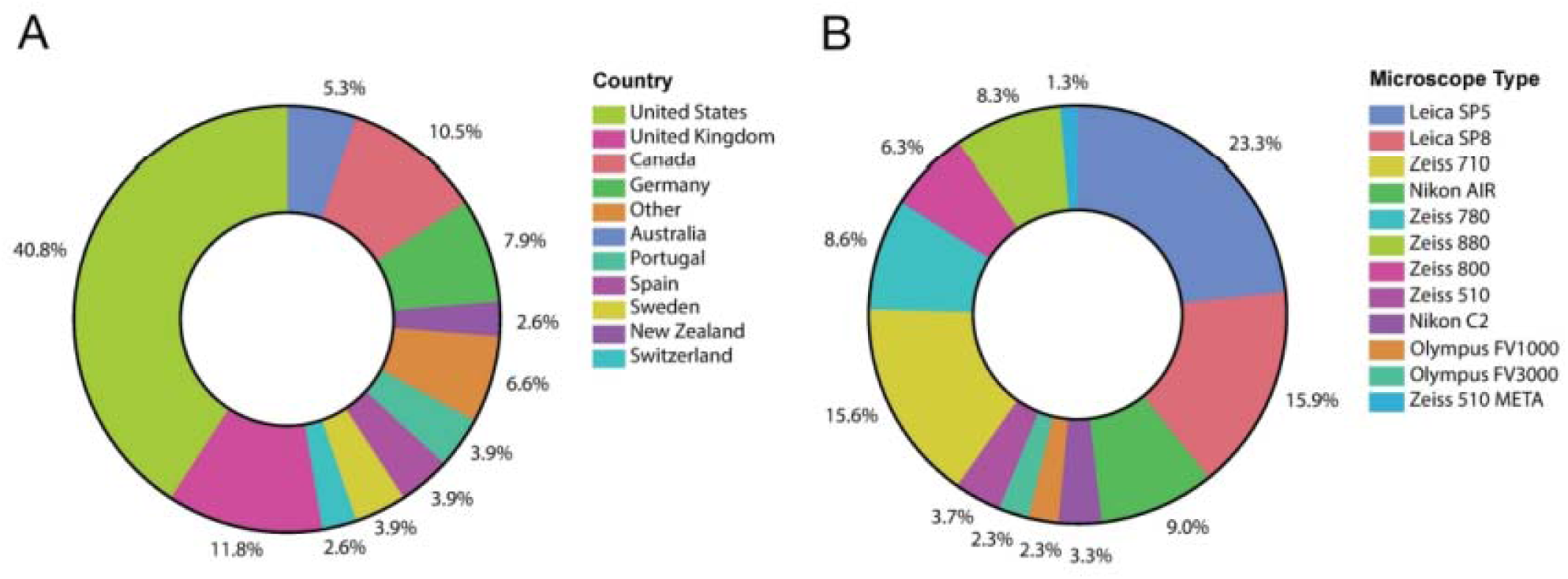
Instruments used during the study and demographics of individual study participants. The study was posted on the confocal list server (http://lists.umn.edu/cgi-bin/wa?A0=confocalmicroscopy) and announced during many in person meetings. Participants were sent sample preparation kits and detailed instructions, protocols and videos for sample preparation and image acquisition. Datasets were sent back to the ABRF-LMRG for analysis. Participants were from more than 12 countries (A) and data was collected using microscopes from all major microscope manufacturers (B). Note that due to the volunteer nature of this group and limited time to work on the project, the data was collected some time ago and data from more recent microscope types is not included.

### Point Spread Function (PSF)/Resolution Analysis

Measuring the PSF of sub-resolution fluorescent microspheres is a reliable method for determining the resolution of a CLSM (Cole, et al., 2011). Study participants prepared 3D samples in CyGel^TM^ containing 1.0 µm diameter orange microspheres for easy focusing on the sample and 100 nm diameter green microspheres to measure the resolution and optical aberrations as a function of imaging depth. Participants were instructed to set up their confocal microscopes to maximize use of the detector dynamic range but ensure there was no saturation. This was to ensure datasets were of high signal-to-noise for robust reproducible resolution measurements. The same imaging settings were used throughout the depth of the sample so microspheres deeper in the sample inherently had lower intensity signals. However, the curve fitting was robust and gave similar resolution values regardless of the amplitude of the microsphere intensity (Supplemental Fig. 4). A total of 140 datasets were received for this part of the study. A subset of 30 datasets passed all criteria for analysis (see Materials and Methods section) and are included in the analysis and figures presented here.

**TABLE 2:** Image Signal-to-Noise-Ratio (SNR) per 2.6-2.7 µm diameter green or red microsphere as a function of depth into the 3D standard sample. Image SNR was measured and presented as explained in the Fig. 4 legend. SNR measurements were grouped by distance from the coverslip into the 3D standard sample by quadrant: quadrant 1 (Q1) closest to the coverslip: 0-25 µm, quadrant 2 (Q2): 25-50 µm, quadrant 3 (Q3): 50-75 µm and quadrant 4 (Q4): 75-100 µm. The Bulk Population mean and standard deviation measurements are given for each NA class over the entire image volume including all depth quadrants.

| Summary of SNR Measurements |  |  |  |  |
| --- | --- | --- | --- | --- |
| NA | Difference in median value Q1:Q4 | Difference in median value as percentage of Max SNR between Q1:Q4 | Bulk Population Mean | Bulk Population StDev |
| 0.7 | 0.08 | -1% | 7.31 | 3.61 |
| 1.2 | -0.86 | 10% | 10.35 | 4.81 |
| 1.25 | 3.17 | -28% | 11.07 | 6.18 |
| 1.3 | 2.98 | -35% | 7.93 | 4.40 |
| 1.4 | 2.67 | -29% | 9.69 | 5.71 |

Changes in the size and shape of the point-spread function (PSF) laterally (x, y) and axially (z) as a function of imaging depth were measured. The lateral full width at half maximum (FWHM_xy_) of the 2D Gaussian fit to the 3D intensity signal from many 100 nm fluorescent microspheres was measured for each depth quadrant in the sample with quadrant 1 being closest to the coverslip (Q1 = 0-24.99 µm, Q2 = 25-49.99 µm, Q3 = 50-74.99 µm, Q4 = 75-100 µm). This gave a measure of the lateral resolution limit of each microscope when imaging this type of 3D sample. Note that the use of the term FWHM_xy_ Mean in the figure refers to the mean value of the FWHM in the x direction and the FWHM in the y direction. However, x and y were not in the same orientation for all microspheres, but x and y were always taken to be perpendicular to one another. The axial resolution of FWHM_z_ was measured in the same way from a 1D Gaussian fit to the 3D intensity signal along the z-axis.

The resolution data was divided into groups of objective lenses sorted by numerical aperture (NA). Resolution data for different NA categories were only included in the analysis and figures if, 1) there were at least 100 independent microspheres detected in the image stack, 2) the PSF was able to be fit (e.g. not a microsphere cluster, see Materials and Methods section for other data pruning criteria) and 3) there were at least three independent datasets for each NA category. Independent datasets may have come from different laboratories but if there was only one high quality dataset for a given NA class, e.g. 0.7 NA air objective lens, then the dataset was not included and is not presented here. The Box and Whisker plots shown in Figure 2 were defined as follows: each box represents all the data points from the 1^st^ to the 3^rd^ quartile, i.e. values between 25%-75% of the median, this defines the interquartile range. The upper and lower whiskers include the values that were within 1.5x the interquartile range. The individual points shown outside of the whiskers are individual PSF measurements that are outside the 1.5x interquartile range. The horizontal black line within the box indicates the median value. The distribution of points within the boxplots is overlaid as a contour plot, the darker the coloration the denser the cluster of data points. For clarity, individual PSF resolution measurements are not shown within the 1.5 interquartile range. Trend lines and a linear fit equation is shown for each NA category (Figure 2).

**Figure 2.**
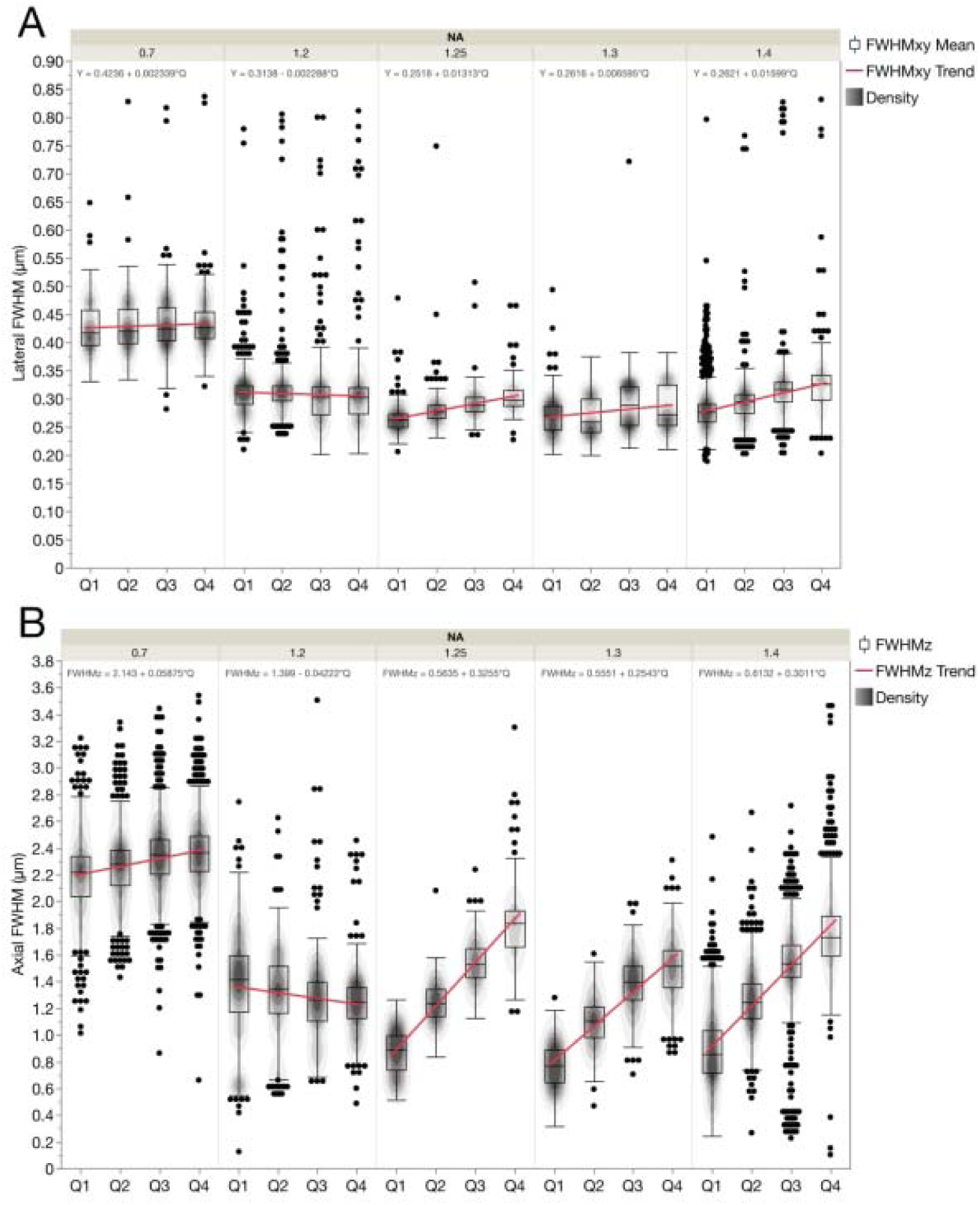
Resolution of microscopes based on PSF curve fitting. Lateral (FWHM_xy_) (A) and axial (FWHM_z_) (B) resolution based on curve fitting of PSFs from image stacks of sub-resolution 100 nm diameter green microspheres as a function of depth in the 3D standard CyGel^TM^ sample. FWHM_xy_ values were measured from 2D fits and FWHM_z_ values from a 1D fit of at least 100 PSFs for each of 30 datasets. Data is shown as a function of depth in four depth quadrants: Quadrant 1 (Q1) closest to the coverslip: 0-24.99 µm, Quadrant 2 (Q2): 25-49.99 µm, Quadrant 3 (Q3): 50-74.99 µm and Quadrant 4 (Q4):75-100 µm. Data was sorted and grouped into categories as a function of objective NA. Box plots show central line that is the population median, upper box is the 3^rd^ quartile, lower box is the 1^st^ quartile and upper and lower whiskers are standard 1.5* the interquartile range (i.e. between 1^st^ and 3^rd^ quartiles). Changes in PSF shape are minimal for lateral PSF fitting (A) and more pronounced with axial PSF fitting (B) due to increased spherical aberrations with depth. Red lines show population trends and the equation for a linear fit to the trend line is shown at the top of each set of boxplots. The contour-line overlays show the relative datapoint density of each plot. The black dots show the outliers. The bulk population mean values, are listed in Table 1 (top) along with standard deviations for these measurements. The n values are the number of microspheres and n = 2724 for 0.7 NA, n = 2984 for 1.2 NA, n = 1145 for 1.25 NA, n = 1146 for 1.3 NA, n = 5941 for 1.4 NA. The n values are the same for Figs. 2 and 3.

When comparing the Q1 data, the lateral resolution (FWHM_xy_) generally increases (smaller FWHM value) with higher NA objective lenses (Fig. 2A). However, the data does have some variability and includes outliers. In general, the mean lateral resolution of the 1.25 NA and above lenses decreased slightly as a function of imaging depth. One contributing factor to this decrease in resolution is that immersion oil objective lenses within the dataset have a significantly higher refractive index (RI) (RI = 1.474 - 1.516) than CyGel^TM^ (RI = 1.365) resulting in spherical aberrations that are more apparent deeper into the sample. The 1.2 NA lenses do not show significant changes in lateral resolution with depth as they are likely water immersion lenses (RI=1.33) that have an RI similar to CyGel^TM^. The lowest resolution 0.7 NA objective lenses showed little change in resolution as a function of depth (Fig. 2A). This is likely due to fact that these lenses have a low lateral resolution (∼400 nm) and large depth-of-field (∼1.0 µm) compared to the 100 nm microspheres making it difficult to detect small changes in lateral resolution (FWHM_xy_) as a function of depth. The impact of depth on FWHM_xy_ could be modeled mathematically, but it is important to point out that the change was very small in most cases (Table 1, top).

Axial resolution (FWHM_z_) measurements gave an indication of the axial resolution of the objective lenses in each NA category as a function of depth in the 3D sample (Fig. 2B). As with the lateral resolution measurements, the FWHM_z_ values were calculated for each NA group and each depth quadrant was compared to theoretical values with Q1 being the quadrant closest to the coverslip (Table 1). Spherical aberrations are expected to have a more significant effect on axial resolution and as expected the FWHM_z_ is more variable with depth (Fig. 2B) than the lateral resolution (FWHM_xy_) (Fig. 2A) for most of the NA categories. As with the lateral resolution, the largest change in resolution with depth is seen with oil immersion lenses where the refractive index of the oil is significantly different than that of CyGel^TM^. The 1.2 NA lenses are mostly water immersion and do not show as much change in axial resolution with depth. Improper correction collar adjustments likely account for the negative slope of the FWHM trend lines for the 1.2 NA objective class (Fig. 2B). Similarly, improper correction collar adjustment of water lenses was a significant factor contributing to artifacts in a past ABRF-LMRG study with about half of lenses showing much higher than expected axial resolution (Cole, et al., 2013). As with the lateral resolution, the 0.7 NA category lenses yielded a small change in the axial FWHM_z_ measurements over depth due to low sensitivity with 100 nm microspheres and high depth of field (Fig. 2B and Table 1, bottom). Resolution data from Fig. 2 is further divided by immersion media and provided in Supplemental Fig. 5. However, note that some immersion media categories did not meet the requirement to have 100 individual data points and 3 independent experiments for PSF determination. It is clear from these plots that the double peaks seen in Fig. 2a and 2b are due to differences in immersion media and lower resolution when there is a large mismatch with the RI of the sample.

In order to further explore the data and see which NA category gave the lowest FWHM values as a population, all measurements from microspheres in all quadrants were merged (i.e. Bulk Population) and plotted. Data was sorted by NA and the lateral (Fig. 3A) and axial (Fig. 3B) resolution values were plotted as one violin plot with data points colour coded by quadrant (Fig. 3; see also, Table 1 “Bulk Population Mean(s)”). The 1.3 NA category shows slightly higher resolution than the 1.4 NA category (Fig. 3A, B), which might have been predicted to perform better based on NA alone. However, this is likely due to the inclusion of glycerol oil objectives in the 1.3 NA class that have an immersion medium refractive index (RI = 1.47) that more closely matches that of CyGel^TM^. There is a clear separation in the distribution of resolution for the microspheres from different depth quadrants that is more apparent at higher NA and with the axial resolution measurements (separation of colors for each quadrant in Fig. 3A, B). Overall the data is of high quality, but there are also some outliers and some distributions demonstrate multiple local maxima in the violin plots. There could be many causes for this including, 1) microsphere aggregates, 2) low signal-to-noise images, 3) poor curve fitting, 4) improper correction collar settings, 5) damaged objective lenses or 6) inhomogeneities in the CyGel^TM^. The data could be explored in more depth for individual microscopes and objective lenses to definitely determine which factors might be contributing.

**Figure 3.**
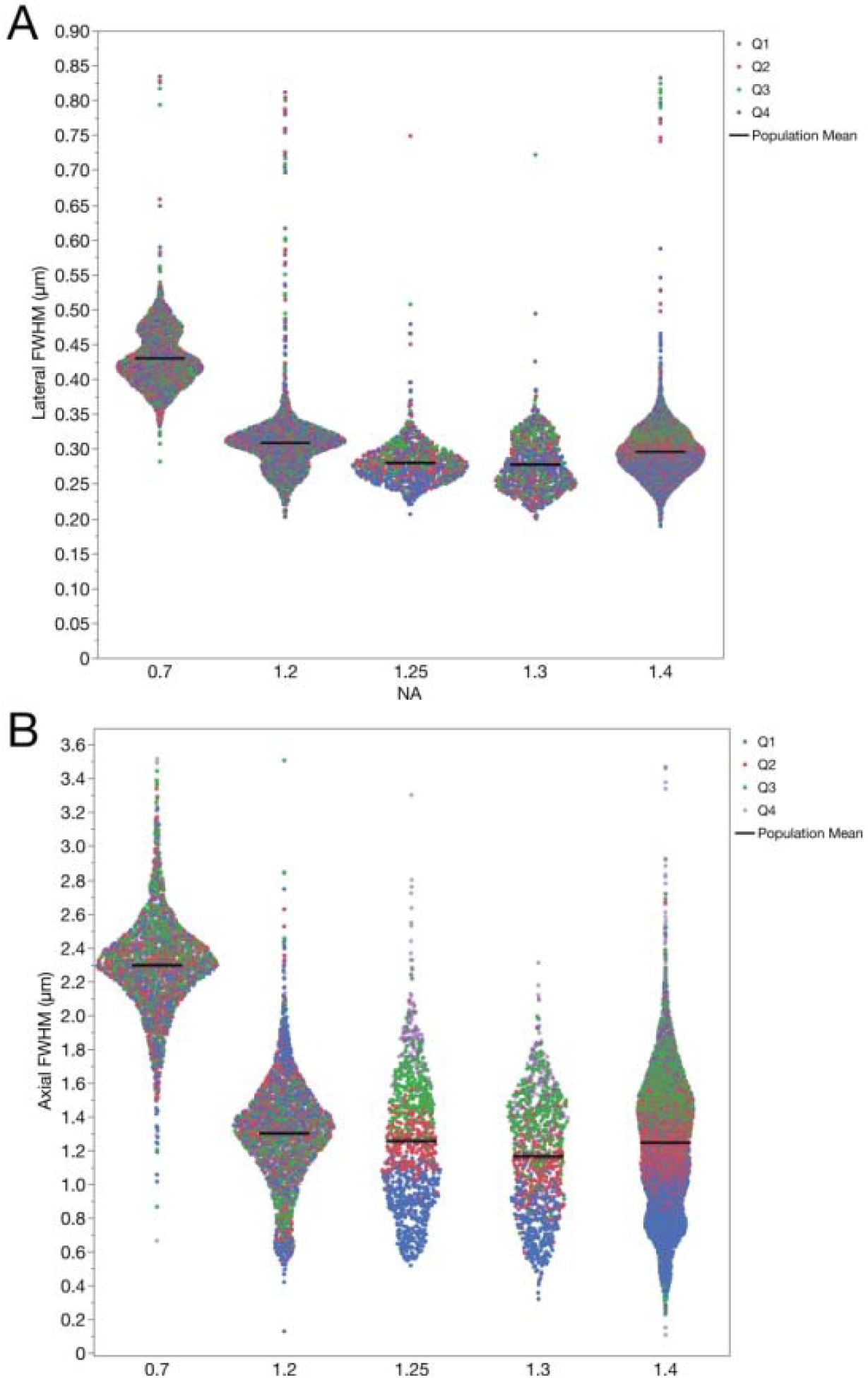
Lateral and axial resolution of the PSFs for data from microspheres at all sample depths as a function of the objective lens NA. Datapoints are plotted for each individual microsphere and color coded by depth quadrant for the lateral FWHM_xy_ (A) and axial FWHM_z_ (B) PSF FWHM resolution. The distribution of the data is displayed as a violin plot with the data organized into groups based on objective lens NA. The increase in the axial PSF FWHM as a function of depth into the sample is evident for the high NA lenses. The black lines mark the bulk population mean values. Summary data is also listed in Table 1 (bottom) along with standard deviations for these measurements.

### Signal-to-Noise Ratio (SNR) Analysis

The SNR was measured from the 100 nm sub-resolution microsphere data by measuring the standard deviation of the difference between the raw intensity data and the fitted Gaussian curve for all x, y data points above the FWHM of the fitted PSF Gaussian functions. Although the number of data points can be small (median of 15 data points above the FWHM), only points for which the correlation between fit and signal was above 90% were chosen. In fact, the majority of the data showed correlation between fit and signal that was close to 98%. Thus, any mismatch is likely attributable to noise rather than bias or misfit. As with the resolution data, data was organized by objective lens NA and depth from the coverslip into the 3D sample by depth quadrants (Fig. 4). As with the trends seen in the PSF data, the SNR was most constant across depth for the 0.7 NA and 1.2 NA lenses and as expected decreased as a function of depth for 1.25 NA and higher lenses. In all cases the SNR was high owing to the high quality of the data sets submitted to the study (Fig. 4, Table 2). There were some outliers, likely due to microsphere aggregates giving high SNRs.

**Figure 4.**
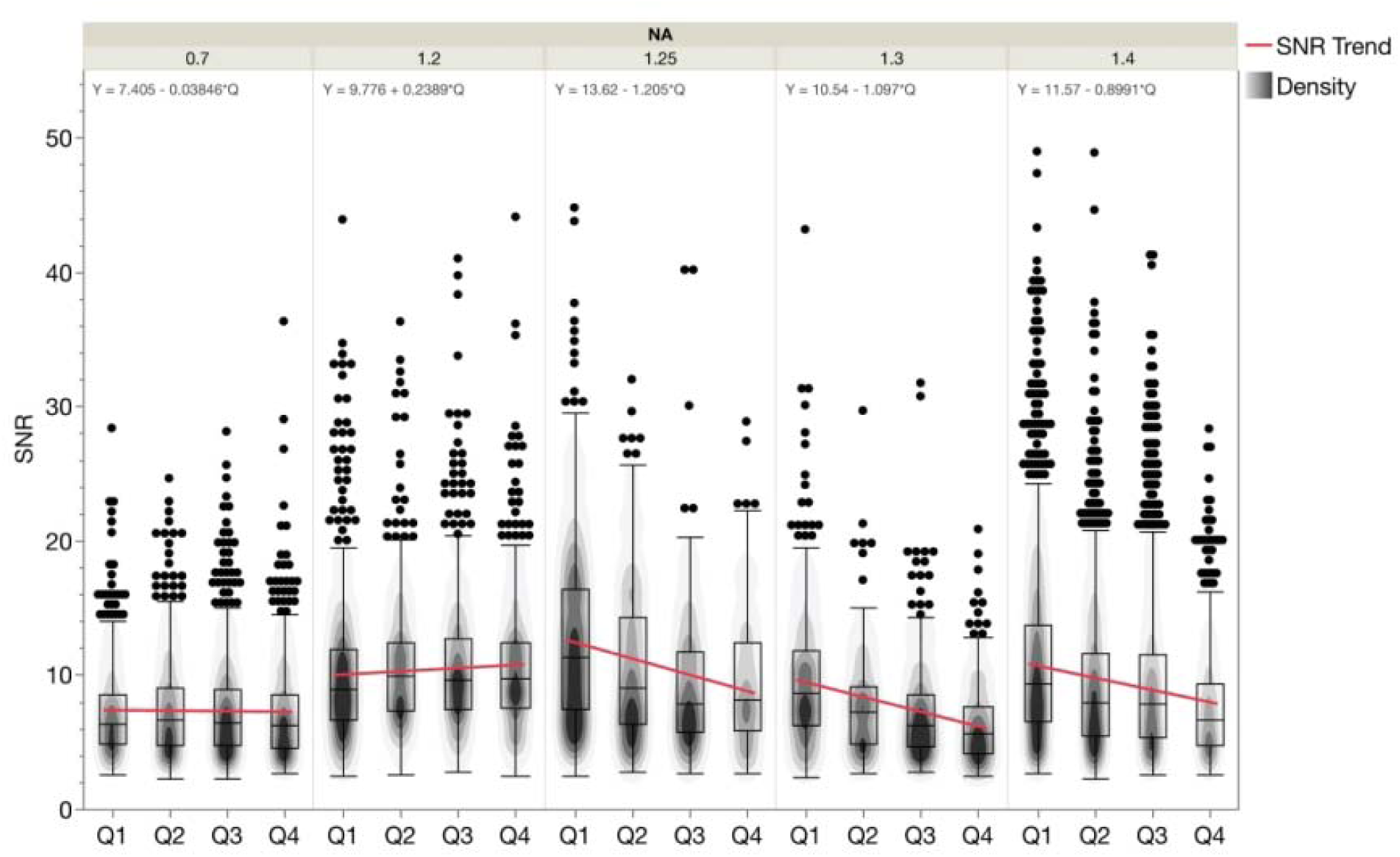
Signal-to-noise-ratio (SNR) as a function of objective lens NA and depth. Box plots show the range of measured SNR values. Data is shown as a function of depth in four depth quadrants: as described in Figure 2 Data was sorted and grouped into categories as a function of objective lens NA. The contour overlays show the data point density for each plot. The trend lines for each NA category are shown in blue with the equation of the line indicated at the top of the plot for each NA category. Boxplot parameters are as described in the manuscript text and the caption for Figure 2. Summary values are provided in Table 2. The n values are the number of microspheres and n = 2722 for 0.7 NA, n = 2981 for 1.2 NA, n = 1145 for 1.25 NA, n = 1146 for 1.3 NA, n = 5937 for 1.4 NA.

### Fluorescence Intensity and Intensity Ratios as a Function of Depth

Study participants prepared samples containing variable intensity green 2.6 µm diameter microspheres with dim (3.7%) or bright (35%) fluorescence intensity signal and red 2.7 µm diameter microspheres with dim (4.7%) or bright (40%) fluorescence intensity signal. For this portion of the study, a total of 191 datasets were received but 91 datasets were eliminated following data pruning (see Materials and Methods). The study participants utilized CLSM instrumentation from Leica, Olympus, Nikon and Zeiss in upright or inverted configurations. Changes in fluorescence intensity as a function of imaging depth were measured on all confocal configurations. A total of 100 datasets passed the validation selection criteria and were used for the analysis. Selection criteria included elimination of datasets if there were saturated intensity values and the requirement for a significant number of microspheres of both dim and bright fluorescence intensity throughout the 3D volume (Fig. 5A). The intensity data were corrected for background by subtracting the background intensity from each voxel and the mean intensity for each microsphere was measured using the spots function in the Imaris software (see Materials and Methods section). As with the resolution data, the results from fluorescent microsphere analysis were separated into four depth quadrants with Q1 near the coverslip, there were losses in intensity with depth and the PSF of the large microspheres became more elongated along the axial direction as a function of depth (Fig. 5B).

**Figure 5.**
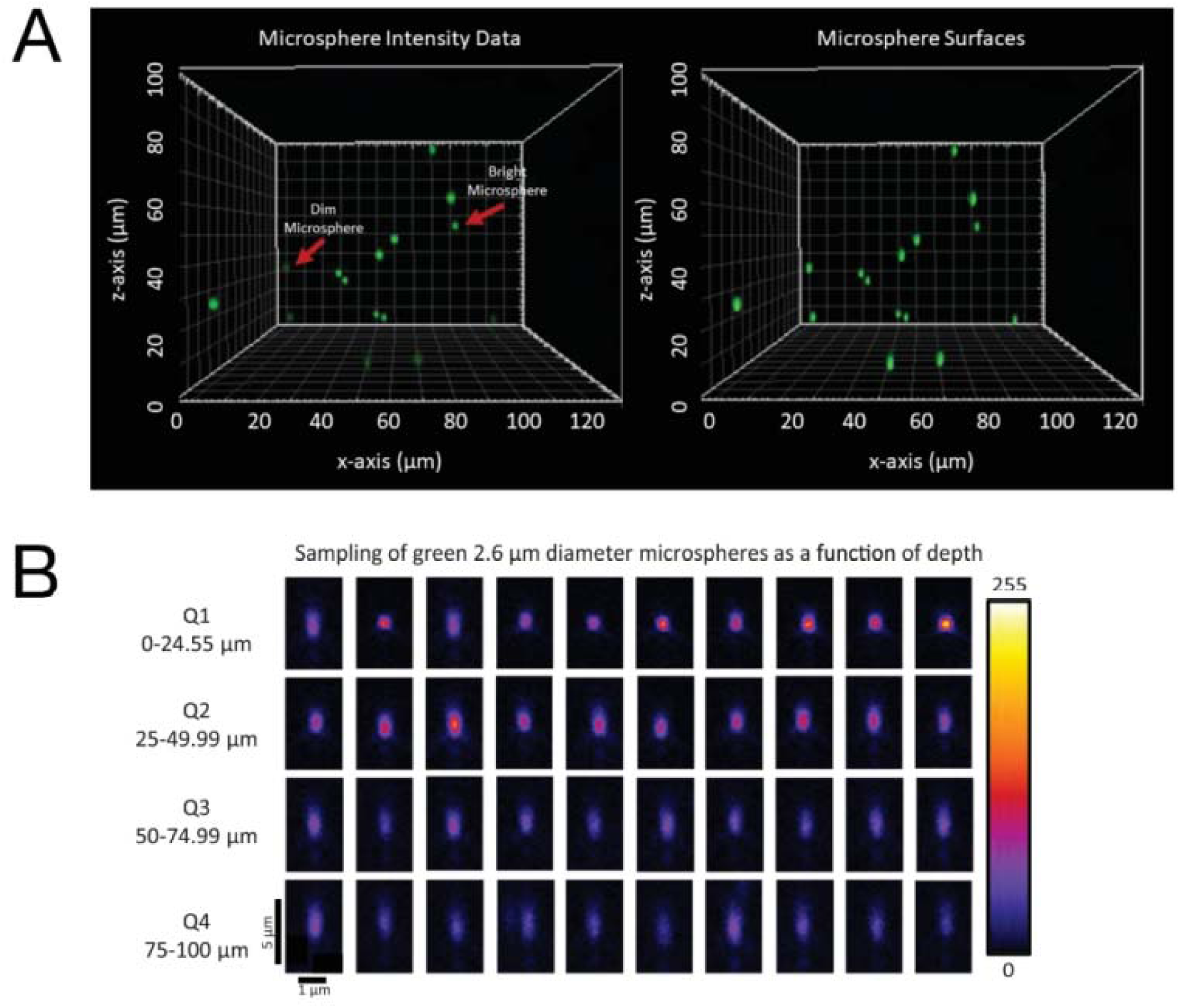
Representative images of 2.6 µm diameter green microspheres as a function of depth. Screen shot from Imaris showing raw intensity (left panel) and Imaris spots surfaces (right panel) for bright (35%) and dim (3.7%) 2.6 µm diameter green microspheres within a 100 µm thick sample of CyGel^TM^ from the xz-axis perspective (A). Spot statistics were measured for every microsphere and exported to an Excel file for further analysis. Representative images of the xz-plane through many different bright green microspheres from different depth quadrants in the 100 µm thick 3D CyGel^TM^ sample showing the axial PSF (B). Quadrant 1 (Q1) closest to the coverslip: 0-24.99 µm, Quadrant 2 (Q2): 25-49.99 µm, Quadrant 3 (Q3): 50-74.99 µm and Quadrant 4 (Q4):75-100 µm. Note the loss of intensity with depth and the elongation of the PSF due to spherical aberrations. Data was collected with a 63x/1/4 NA oil immersion objective lens on a Zeiss LSM800. The 8-bit intensity scale is indicated on the right side.

Absolute fluorescence intensity is not consistent across different CLSMs. Variables such as laser power, objective light transmission efficiency, NA, detector sensitivity, and optical components in the light path make it impossible to directly compare fluorescence intensity values, regardless of the microscope manufacturer. To compare microsphere intensities across samples and depths within the 3D sample, a method to normalize the data and compare relative values was developed. Within each dataset, the intensity of each microsphere was normalized to the intensity of the brightest microsphere in the first quadrant (Q1). Thus, datasets that had no microspheres in the first quadrant were eliminated from further analysis. The normalized intensities were then averaged over all microspheres in each quadrant for the dataset.

The data were divided into subsets from objectives with different NA ranges including 0.7 NA – 0.75 NA air and oil immersion lenses (Fig. 6A), 0.8 NA – 0.85 NA air or water lenses (Fig. 6B), 1.2 NA water or 1.25 NA oil immersion lenses (Fig. 6C) and 1.3 NA glycerol, 1.3 NA oil or 1.4 NA oil immersion lenses (Fig. 6D). The wide range of normalized intensity values is due to different immersion medium, microsphere intensity, the high dependence on the normalized intensity of the intensity of the brightest microsphere in Q1 for each data set and the inclusion of microspheres from the entire 25 μm depth for each quadrant (Fig. 6). This is apparent where losses in fluorescent intensity were less significant with better IR matching. For example, the losses in intensity with depth are lower with the 0.7 NA glycerol immersion lens (Fig. 6A), the 0.8 NA water lens (Fig. 6B, Supplemental Fig. 1A) and the 1.2 NA water immersion lens (Fig. 6C, Supplemental Fig. 1B). In turn, there is little difference between the 1.3 and 1.4 NA oil immersion lenses (Fig. 6D). Thus, this indicates that the loss in intensity is due to spherical aberrations caused by refractive index mismatches between the objective immersion medium and the CyGel^TM^ plus scattering of emission light as it travels back through the gel to be detected by the objective lens.

**Figure 6.**
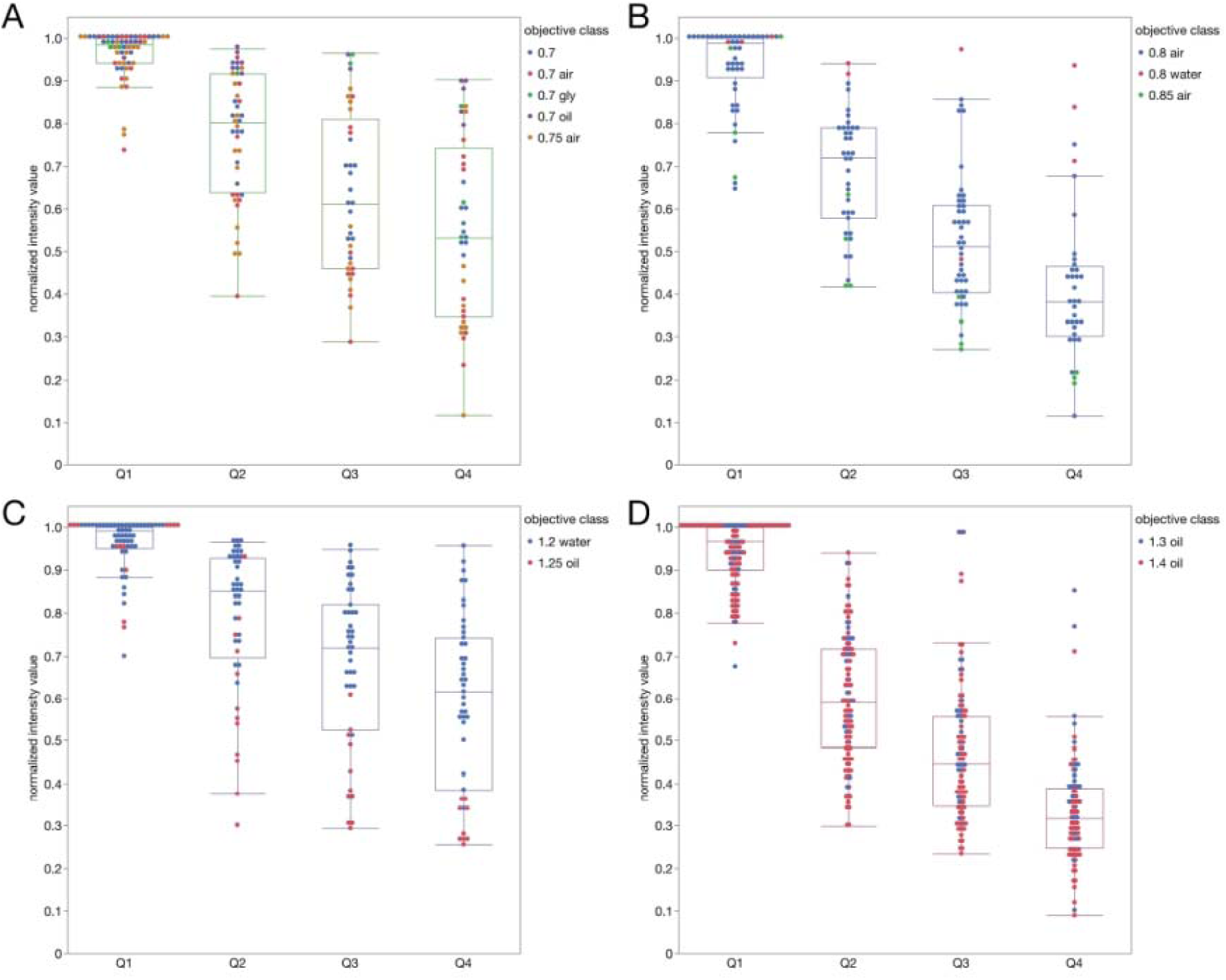
Relative fluorescence intensity as a function of depth for all bright green and red fluorescent microspheres. Relative intensity data for 2.6 µm diameter bright green and 2.7 µm diameter bright red microspheres as a function of depth into the 3D gel sample. Intensities were normalized for each dataset relative to the brightest microsphere in Q1. Data is a compilation of 81 datasets, collected on 41 different CLSMs. Data is sorted by NA = 0.7-0.75 (A), NA = 0.8-0.85 (B), NA = 1.2-1.25 (C) and NA = 1.3-1.4 (D). Each data point is from a single microsphere and is color coded based on the objective NA and immersion medium. The n values are the number of microspheres and n = 176 data points for panel A, n = 168 data points for panel B, n = 195 data points for panel C, n = 453 data points for panel D.

Having access to microspheres of a known ratio of intensities within a 3D sample allowed for the calculation of the ratio of the intensity of the bright/dim green (Fig. 7A) or red (Fig. 7B) microspheres as a function of depth. After pruning the data, results from 40 different CLSMs with objective lens magnifications ranging from 20X to 63X, and NA ranging from 0.7 to 1.40 were included in the analysis. Importantly, separating the ratio data as a function of NA of the different lenses revealed that there was no dependency of the ratio on NA, immersion media or quadrant within the 3D sample (Supplemental Figs. 2 and 3). The fact that the same ratio was measured regardless of objective lens NA, immersion medium, type of CLSM, normalization, microscope operator clearly demonstrates that this is a robust standard sample.

**Figure 7.**
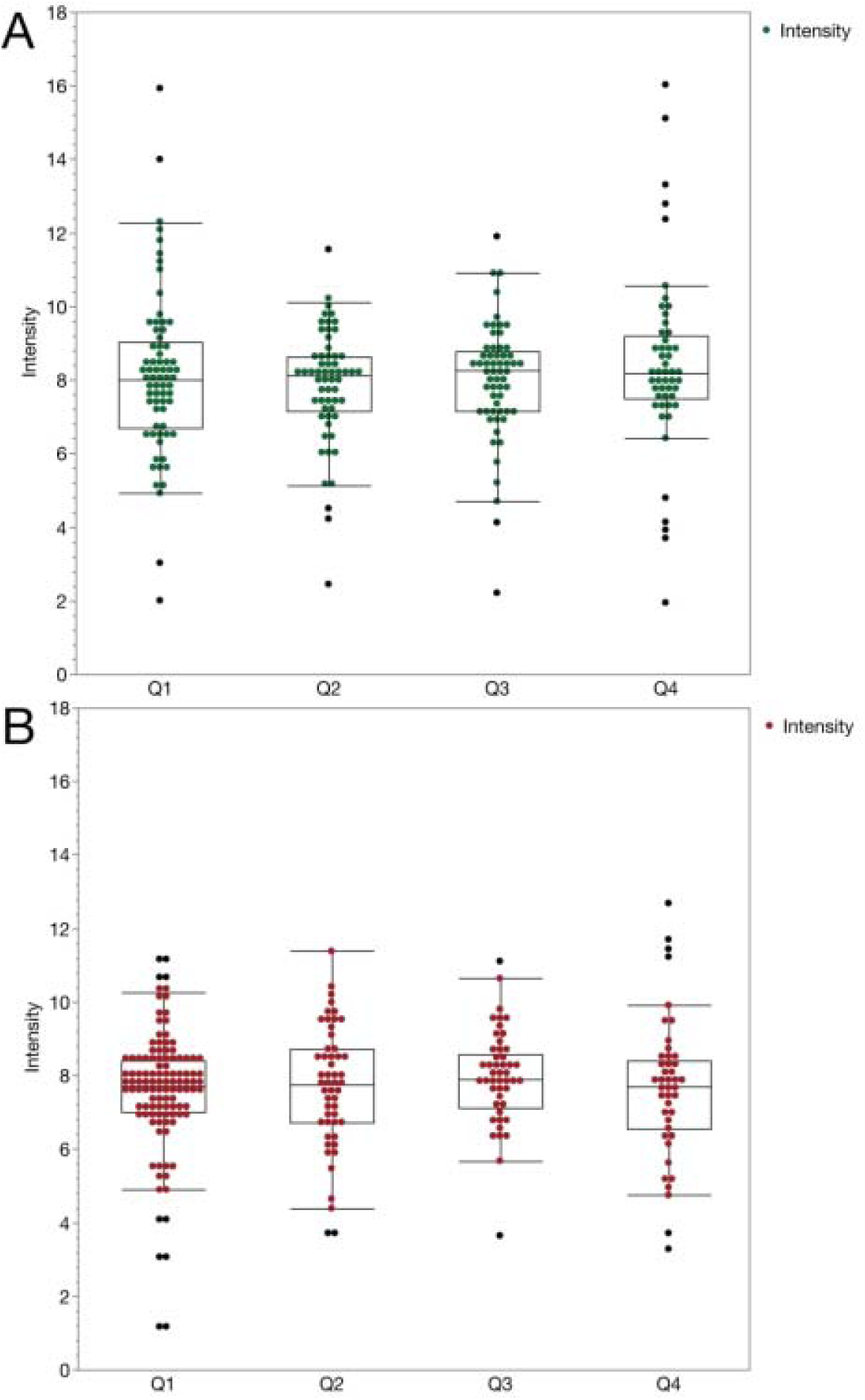
Bright over dim fluorescence intensity ratios for green and red fluorescent microspheres as a function of depth. Fluorescence intensity ratios (Bright/Dim) for 2.6 µm diameter green microspheres (A) and 2.7 µm diameter red microspheres (B) as a function of depth in the 3D CyGel^TM^ sample. Data is a compilation of 100 datasets, collected on 47 different CLSMs. Objective lens magnification ranged from 20x to 63x and NA from 0.7 to 1.4. The depth quadrants and Boxplot parameters are as in Figure 2 caption. Spots show outliers. The n values are the number of ratios measured and n = 237 measurements for the green ratio data. n = 254 measurements for the red ratio data.

Overall, the mean ratio for the bright/dim green microspheres was 8.08 (Table 3). This is significantly different from the ratio determined by ThermoFisher for this lot of microspheres which was 8.5 (40%/4.7%). Essentially, the ratio obtained through our analysis was 5% lower than the expected value. The mean ratio of the bright/dim red microspheres was 7.68 (Table 3) while the expected ratio was 35%/3.7% or 9.6. Thus the measured ratio was 20% lower than the expected value. As mentioned above, these measured values were also consistent across all depths, all instruments, immersion medium and objective lens types tested (Supplemental Figs. 2 and 3).

**TABLE 3:** Summary of Bead Intensity Ratio Measurements. This table gives the mean, median and standard deviation for the intensity ratio calculations for each depth quadrant for the green microspheres and the red microspheres. The Bulk Population column gives mean, median and standard deviation measurements for each color bead over the entire image volume including all depth quadrants.

| Bead Intensity Ratio Measurements |  |  |  |  |  |  |
| --- | --- | --- | --- | --- | --- | --- |
|  | Bead Color | Q1 | Q2 | Q3 | Q4 | Bulk Population |
| Mean | Green | 8.02 | 7.83 | 8.03 | 8.52 | 8.08 |
| Median | Green | 8.00 | 8.11 | 8.31 | 8.20 | 8.12 |
| StDev | Green | 2.28 | 1.53 | 1.60 | 2.62 | 2.04 |
| Mean | Red | 7.57 | 7.69 | 7.94 | 7.65 | 7.68 |
| Median | Red | 7.74 | 7.75 | 7.89 | 7.72 | 7.77 |
| StDev | Red | 1.67 | 1.65 | 1.29 | 1.94 | 1.65 |

Lower ratios could be due to saturation of fluorescence and/or fluorophore quenching of bright microspheres leading to a numerator in the ratio that is of lower intensity than expected. It is also possible with the manual adjustment of the threshold settings in the Imaris software for the dim microspheres that the size was sometimes underestimated. This would miss dimmer intensity pixels near the edges of the microspheres thus increasing the mean microsphere intensity in the denominator and decreasing the intensity ratio. Finally, z-axis sampling was at 1 µm so the position of the microsphere relative to the image planes could vary throughout the sample. If one of the image planes is not centered around the middle of the microsphere this could result in an underestimate of the microsphere intensity.

## DISCUSSION

This study presents a detailed list of reagents and materials, protocols and analysis methods to measure lateral and axial resolution, optical aberrations, SNR, intensity loss and relative intensity all as a function of depth in a 3D standard sample that is a suitable tissue mimic of 100 µm depth. The cost of preparing such samples is reasonable, the protocols and analyses are straightforward, and they should provide the bioimaging community with valuable tools towards rigor and reproducibility in quantitative light microscopy (Lee & Kitaoka, 2018; National Academies of Sciences, 2019). The standards can be used to establish baseline instrument performance, track performance over time, identify instrument issues, build standard datasets for image corrections and add to the toolbox required for high quality quantitative bioimaging. Importantly, these standards provide a 3D tool, of which few exist in the field, to mimic tissue samples and make standard measurements up to 100 µm into the sample to complement more readily available tools that test performance at or close to the coverslip.

Lateral and axial resolution measurements from 100 nm diameter sub-resolution microspheres gave somewhat expected results. The measured resolution limits were higher than predicted by theory as is often the case with experimental systems (Cole, et al., 2011). “There could be several reasons why this is true. The most significant reason is refractive index mis-match effects and spherical aberration as the microspheres are further from the coverslip. Even within Q1 the depth range is 25 µm. This underperformance phenomenon is also described in Hell et al. (1993), Visser and Oud (1994) and in Sheppard and Torok (1997) texts, which describe refractive effects. Visser points out that the distance that the light is traveling through the oil is changing as one focuses into the sample, this changes the focal point and enlarges the apparent volume of the object. The apparent degree to which this occurs is heavily dependent on the analysis method used to study the effect (Besseling, et al., 2015). Another potential source of underperformance is that the instruments have not been recently quality control checked and calibrated. Thus it is possible that several smaller performance- reducing issues limit the resolution that can be realized. This points to the importance of using standard samples and protocols such as the ones presented here to benchmark each microscope and lens on a regular basis and identify when repairs and maintenance are needed.”

The FWHM values determined had a broad distribution both within and between datasets, likely partially due to some microsphere aggregates in the sample. Additional filtering for lower intensity PSF peaks could be implemented to mitigate this contribution. Resolution decreased as a function of depth and emphasized the importance of refractive index matching of immersion medium and sample mounting medium especially when imaging in the depth quadrants that were more than 25 µm away from the coverslip. Sample preparation and image acquisition protocols for measuring resolution from the PSF in 3D are presented here and, importantly, MATLAB software was developed to streamline the image analysis. The MATLAB PSF code is available on the ABRF-LMRG GitHub page and can be used to automatically analyze and output PSF data for the lateral and axial FWHM for hundreds of microspheres in single or multiple datasets. The importance of measuring many data points for a single microscope with a specific objective lens and image acquisition settings is evident by the high variability within the data. Future tools could expand the capabilities of the analysis such as automation of data outputs to report resolution as a function of imaging depth and radial location relative to the center of the field-of-view. This type of more detailed analysis would be useful in testing new equipment, new microscope lenses and for ongoing evaluation of instrument performance. The SNR data followed a similar trend to the resolution with lower SNR deeper into the sample and also when there was index of refraction mismatch. This trend was dominated by the loss of signal with depth due to spherical aberrations and light scattering.

The variable intensity microspheres provide an excellent tool to assess sensitivity, relative intensity, intensity decay with depth into a sample and SNR. Our results show how changes in intensity as a function of depth can be measured. The bright and dim, green or red microspheres all give similar results. This means that changes in intensity over depth can be measured using 3D samples such as these microspheres in CyGel^TM^ or in other matrices to calibrate intensity loss in specific biomaterials. The resulting standard curves can then be used to calibrate and correct intensity data for loss as a function of depth due to phenomena such as light scattering or spherical aberrations. *<u>The data also point out how important it is not to quantitatively</u> <u>compare intensities at different depths when imaging deep into samples unless the intensity loss is calibrated and corrected before intensity measurements are made.</u>* <u>Fortunately, intensities at the same or similar depth into the sample can be compared as relative values do not change as a function of depth.</u>

Interestingly, for both the green and red microspheres, the relative intensity values measured in this study were lower than expected based on the product specifications. There is high confidence in the values measured here as they come from many different labs, microscopes, instrument settings and objective lenses and they were handled according to manufacturer recommendations. This is a robust standard sample and intensity ratios were reproducible and comparable across many different objective lenses, CLSM platforms, immersion media with dozens of different researchers preparing and imaging the samples. Thus, the chance of the measured values being the same across many laboratories and many instruments due to the same or similar technical errors would be extremely low. Other possible reasons for the low ratios could be due to fluorophore saturation or quenching of the fluorescence in the bright microspheres due to high fluorophore density. This was previously seen with the 100% InSpeck^TM^ Green microspheres (Lee, et al., 2014). Along these same lines perhaps the bright microspheres have a higher relative photobleaching during the high resolution CLSM image collection or have a higher relative quenching of fluorophores at the microsphere surface due to interactions with the CyGel^TM^ media. It is also possible that the thresholding of the dim microspheres only selected the bright core overestimating their mean intensity. To increase the precision of this assay, five of the intensity standard microspheres in the InSpeck^TM^ kits could be used to generate a full standard curve of intensity as a function of depth into the standard sample. However, as long as the same microspheres are used over time to monitor instrument performance and intensity ratios are used then the absolute intensity ratio values are not critical.

## CONCLUSIONS & FUTURE DIRECTIONS

It is our hope that this study presents researchers, particularly imaging scientists in core facilities, with some new tools to add to their toolbox for rigor and reproducibility of quantitative fluorescence bioimaging. Great care was taken in determining the ideal type of sample, sample medium and developing detailed protocols. Protocols are available for sample preparation including videos, for quantitative image acquisition and for image analysis. The MATLAB code for PSF identification and fitting is available for the community to download and use. The CyGel^TM^ samples can be kept for 1-2 weeks at room temperature but do dry out over time. Fortunately, new standard samples can be made in a few minutes once all the materials are available.

It would be ideal to have a mounting medium that could be tuned for the refractive index to match different biological samples and that once hardened could be kept and stored for years at a time. The standard sample could relatively easily be adapted for different mounting media including media with different refractive indices and media that are stable at 4°C. The microspheres and the image acquisition protocols would not need to be adjusted and the sample preparation would only need minimal modifications. Ideally, these microsphere samples could also be included in 3D samples with the specimen that will be imaged. That way resolution, intensity and aberrations within the specimen under study can be measured and corrected for.

Another major hurdle is the need for automated software tools to facilitate microscope quality management. Ideally automated tools for image acquisition, image analysis and data analytics could be integrated into the microscope image acquisition software and could provide a daily report of the instrument status after 2D and 3D standard samples are placed on the microscope and imaged. These reports could then provide longitudinal data about each instrument, be stored digitally, archived and shared across the community. This would provide imaging facilities, microscope manufacturers, reviewers, journals and granting agencies with the tools they need to ensure rigor and reproducibility in quantitative bioimaging. The standard samples and protocols presented here are one step along the bioimaging communities’ path to this ideal future.

The work of the ABRF-LMRG and the more recently established international QUAREP-LiMi group are putting microscopy standards at the forefront. Community engagement and a focus on providing community agreed upon standards, protocols and tools for image analysis will enable vast improvements in reproducibility in light microscopy. The sheer size of the QUAREP-LiMi community along with efforts to engage corporate partners including microscopy manufacturers, manufacturers of microscope accessories (e.g. light sources, cameras), reagent companies (e.g. the corporate partners on this study, BioStatus Inc, ThermoFisher) and other key stakeholders will ensure we continue to improve microscope quality management.

For example, another challenging aspect of fluorescence imaging is the temporal instability of lamps or laser-based light sources. This instability directly affects the intensity values recorded within images and thus negatively impacts quantitative comparisons of fluorescence intensity within single image data sets (3D or timelapse) and images collected over several imaging sessions that may be collected days, weeks or months apart (Jonkman, et al., 2020; Mubaid, et al., 2019; Stack, et al., 2011). QUAREP is working towards improving this situation with the first QUAREP-LiMi community developed protocol for “*Illumination Power and Stability”* that has recently been published on protocols.io (https://www.protocols.io/view/illumination-power-and-illumination-stability-5jyl853ndl2w/v1). The protocol is available and open for comments so it can continuously be developed and improved with the global community. Similarly, BioImaging North America (BINA) has started a pilot project, modeled after the French program developed by Microscopie de Fluorescence Multidimensionnelle (MFM), where a microscopy quality control kit (i.e. metrology suitcase) and protocols will be shipping throughout North America to enable microscopists of all levels of expertise to quality control their microscopes. The suitcase includes a power meter for measuring illumination power and stability. Several other groups have developed devices to measure incident light power stability over time combined with a cell phone application (Dormann, 2019) or using an objective turret mounted device (Grunwald, et al., 2008) and could be included in future versions of the suitcase. A commercial solution from Argolight includes a microscope slide with a built-in power meter and accompanying software for monitoring power (ArgoPowerHM, Argolight, Pessac, France). The suitcase also includes a TetraSpeck^TM^ 4-color microsphere slide and a PSFCheck slide for measuring microscope resolution.

Finally, the ABRF-LMRG is running a fourth microscopy quality management study on the reproducibility of image analysis. The concept of the study is that all participants will gain access to the same well characterized computer-generated synthetic images, segment the objects and make quantitative measurements. Image analysis methods and software will be diverse but everything will be shared. Analysis results will be pooled and synthesized to see global results and understand aspects of reproducibility in image analysis and identify potential sources of errors and artifacts. For more information and to get involved see https://sites.google.com/view/lmrg-image-analysis-study.

Overall, we are confident that this work provides the community with a new and robust 3D tissue-like standard and that together with the global community we are well on our way towards continuous improvement of rigor and reproducibility in light microscopy (Boehm, et al., 2021).

## Author Contribution Statement in alphabetical order

**Benjamin Abrams**, development and beta-testing of multiple rounds of samples. Writing and refining the acquisition protocol for one of the target instruments. Key role in helping to develop a system for collecting the microscope metadata for image analysis and in helping to develop the initial image analysis workflow for the PSF portion of the study. Helping to analyze the PSF data. Helping to write, create figures for and edit the manuscript.

**Carol J. Bayles,** Tested early samples for feasibility. Using the final sample collected data on two microscope systems and submitted for the study.

**Claire M. Brown**, early development and testing of several rounds of sample development. Key role in development of the final sample, oversight of the project, development and oversight of sample preparation protocols, development of methods for signal and noise measurements in 3D datasets in Imaris, writing and development of analysis protocols, presentation of data at ABRF annual meetings, writing and editing of the manuscript.

**Linda M. Callahan**, development and testing of multiple rounds of samples. Writing and refining the image acquisition protocol for one of the confocal microscopes used for this study. Involved in the PSF data initial imaging protocols and image analysis development. Assisted with writing and editing of the manuscript. Participation in LMRG virtual meetings for ABRF Study #3 development, deployment, and analysis.

**Lisa A. Cameron,** participated in early discussions of study design and methodology, virtual LMRG meetings, aided troubleshooting of launch methodology.

**Richard Cole**, conceptualized the idea for the project based on the need for an affordable easy to prepare and measure 3D microscopy standard. Early development and testing of several rounds of sample development. Key role in development of the final sample, preparation, protocols and analysis.

**Rebecca Deagle,** development and testing of several rounds of sample development. Key role in the development of final sample kit design, preparation protocols, and data analysis. Key role in the development of analysis protocols for SNR measurements in 3D datasets in Imaris. Creating and distributing the instructional video for sample preparation. Maintenance of LMRG meeting minutes and presentation preparation for ABRF annual meetings.

**Anne-Marie Girard**, early testing of several rounds of sample type and preparation including duckweed, nanofibers and CyGel^TM^. Early testing of protocols on Zeiss LSM 510 and Zeiss LSM 780 prior to study samples being sent to participants.

**Kristopher E. Kubow,** contributed to discussions of study design, methodology, and analysis. Helped test and troubleshoot the sample preparation and imaging protocols. Presented data at ABRF annual meetings.

**Thomas Pengo,** participated in discussions regarding PSF analysis, developed and performed the image analysis of the collected PSF data, contributed to the statistical assessments and final analysis, and helped with the corresponding sections of the manuscript.

**Joshua Z. Rappoport**, participated in LMRG virtual meetings, early discussions of study design and methodology, aided troubleshooting of launch methodology, and contributed to manuscript writing and editing.

**Megan Smith** headed communications with study participants and managed sample distributions to participants. Compiled, vetted, and analyzed raw image data as well as statistical data. Writing and editing of the manuscript.

**Nelly Vuillemin**, image analysis and rendering of results for the Signal to Noise Ratio and Sensitivity on imaging depth parts. Development and testing of protocols. Realization of figures to present statistical results. Participation in numerous discussions and troubleshooting discussions. Helping to write and edit the manuscript.

**Tse-Luen Wee,** development and testing of several rounds of sample development. Key role in development of final sample kit design, preparation, and protocols. Execute and oversight of the project, development of sample preparation and imaging protocols, presentation of data at ABRF annual meetings, writing and editing of the manuscript.

## ACKNOWLEDGMENTS

Thank you to the ABRF for funding and continued support in developing light microscopy standards and protocols. The authors would like to acknowledge significant support in the form of human resources and computing resources from the Advance BioImaging Facility (ABIF) at McGill University. Thank you to Molecular Probes/ThermoFisher and the University of Minnesota shared multi-scale microscopy facility for providing fluorescent microsphere samples and BioStatus Ltd. for providing CyGEL^TM^ and covering the cost and logistics of shipping the sample kits to the international study participants. Thank you to Kary L. Oakleaf for consumables selection for 3D sample matrix and early testing of sample preparations. Thank you to all of the study participants: Pablo Ariel, Constadina Arvanitis, Laszlo Barna, Carol Bayles, Nicole Bouffard, Sally Boxall, Michael Cammer, Heather Cartwright, Patrizia Casalini, James Chambers, Wai Chan, Elodie Chatre, Crystal Chaw, Nicholas Condon, Rebecca Deagle, Steffen Dietzel, Joe Dragavon, Judith Drazba, Esteban Fernandez, Julia Fernandez-Rodriguez, Claudia Florindo, Katherina Garcia, Cathy Gillespie, Maria Gomez Lazaro, Nicola Green, Hella Hartmann, Joachim Hehl, Kristina Jahn, Hannah Johnson, James Jonkman, Paivi Jordon, Alexander Jurkevich, Noriko Kane-Goldsmith, David KIrchenbüchler, Elke Küster-Schöck, Vibor Laketa, Chris Law, Sylvie Le Guyader, Stephen Lentz, Julie-Christine Levesque, Kevin Mackenzie, Karen Martin, Arvydas Matiukas, Joseph Mazurkiewicz, Ewan McGhee, Bruce Mockett, Dale Moulding, Jonathan Mulholland, Tobias Nyberg, Ken Orndorff, Scott Page, Ana Paula Rodrigues, Marc Reinig, Paul Reynolds, Juan Luis Ribas, Jaqueline Ross, Craig Russell, Rebecca Saleeb, Paula Sampaio, Ana Santos, Britta Schroth-Diaz, Tobias Schwarz, Joel Sheffield, Alan Siegel, Martin Stoeckl, Andrea Stout, Xuejun Sun, Sarah Swanson, Darren Thomson, Chloe van Oostende-Triplet, Jinsong Wang, Michael Weis, Silke White, Tong Yan, Christopher Yip, Zbigniew Mikulski.

MATLAB code is available at https://github.com/orgs/ABRFLMRG/

---

**Supplemental Figure 1.**
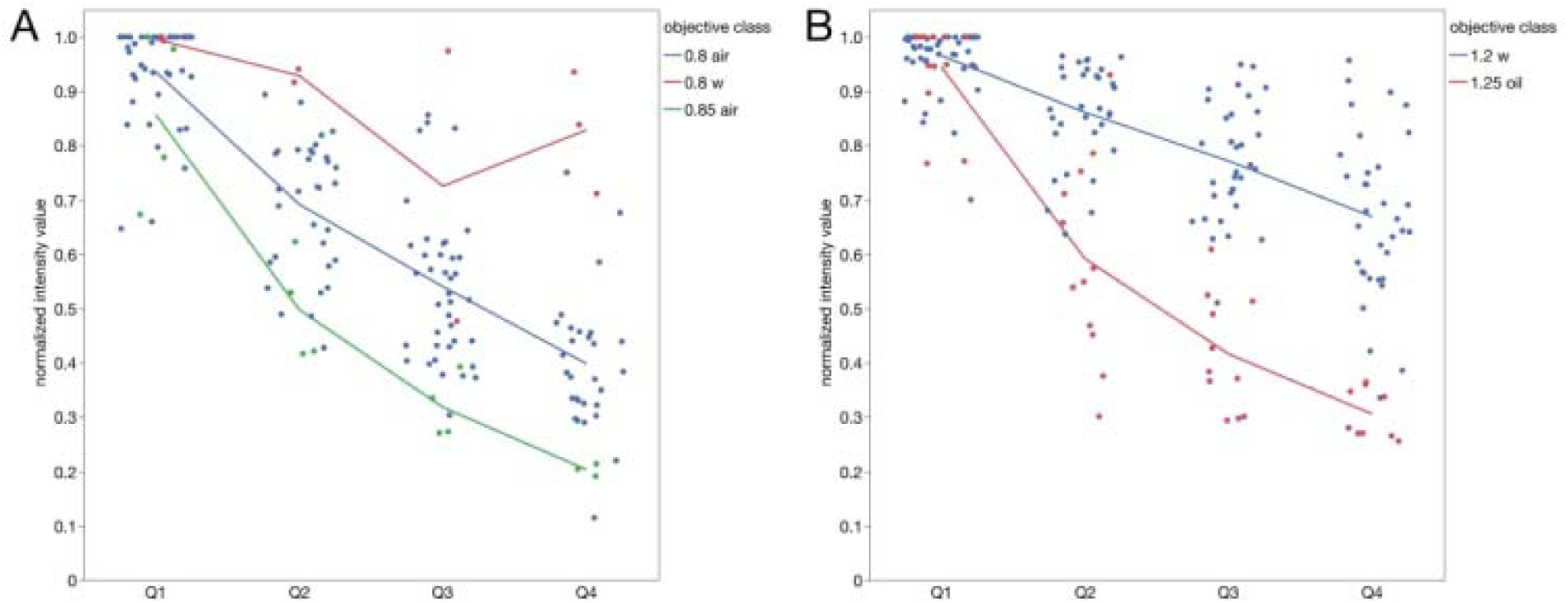
Alternative plotting for two of the NA categories shown in Figure 6. Line graphs plot are broken out for individual objective categories showing the intensity versus depth performance for the 0.8 air, 0.8 water and 0.85 air objectives (A) and for the 1.2 water and 1.25 oil objectives (B).

**Supplemental Figure 2.**
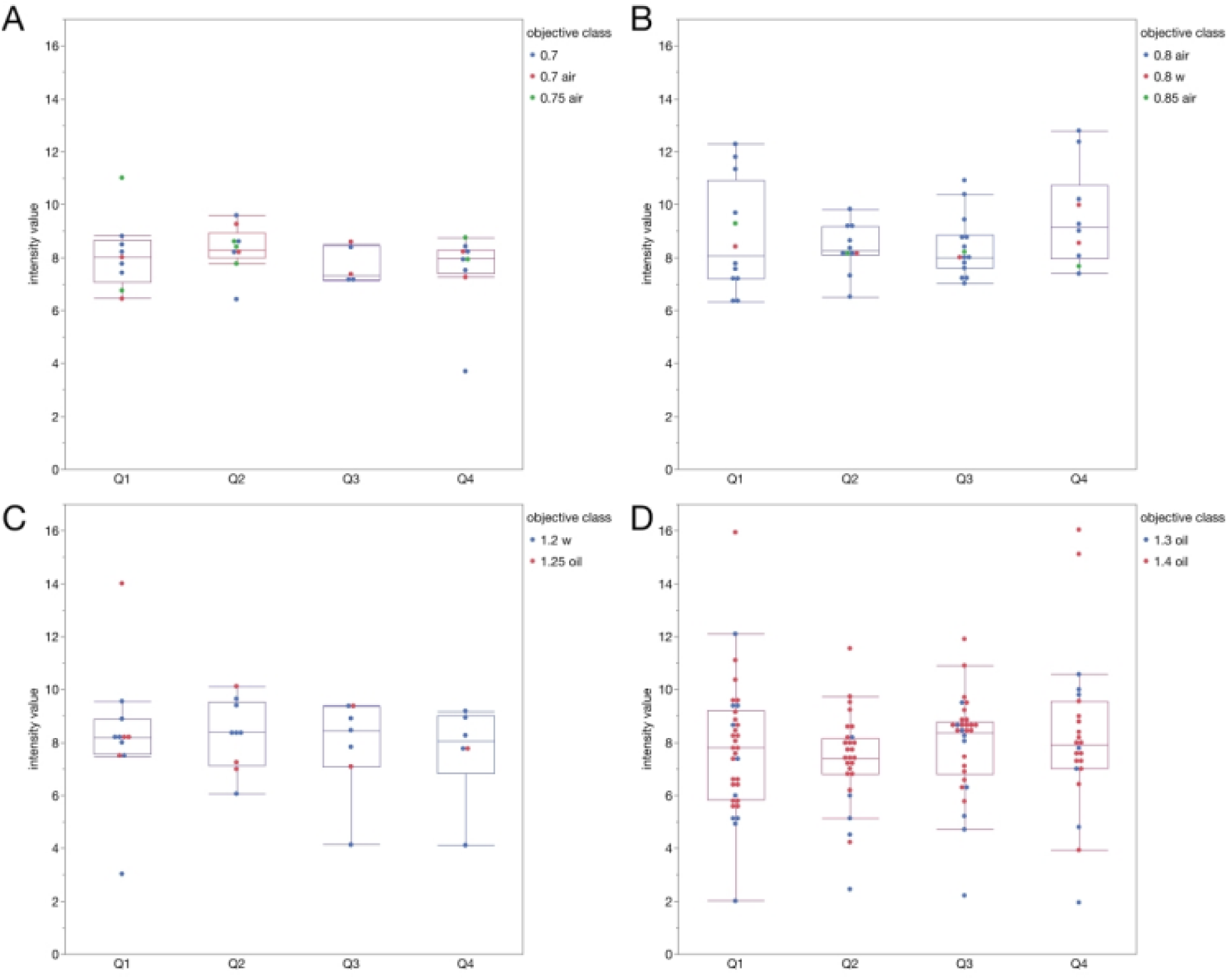
Intensity ratio data broken out by NA. This plot shows the intensity ratio plots for the green microspheres broken out by depth and NA group. 0.7, 0.7 air and 0.75 air (A) 0.8 air (B) 1.2 water, 1.25 oil (C) 1.3 oil, 1.4 oil (D). The ratios are essentially unaffected by depth or NA.

**Supplemental Figure 3.**
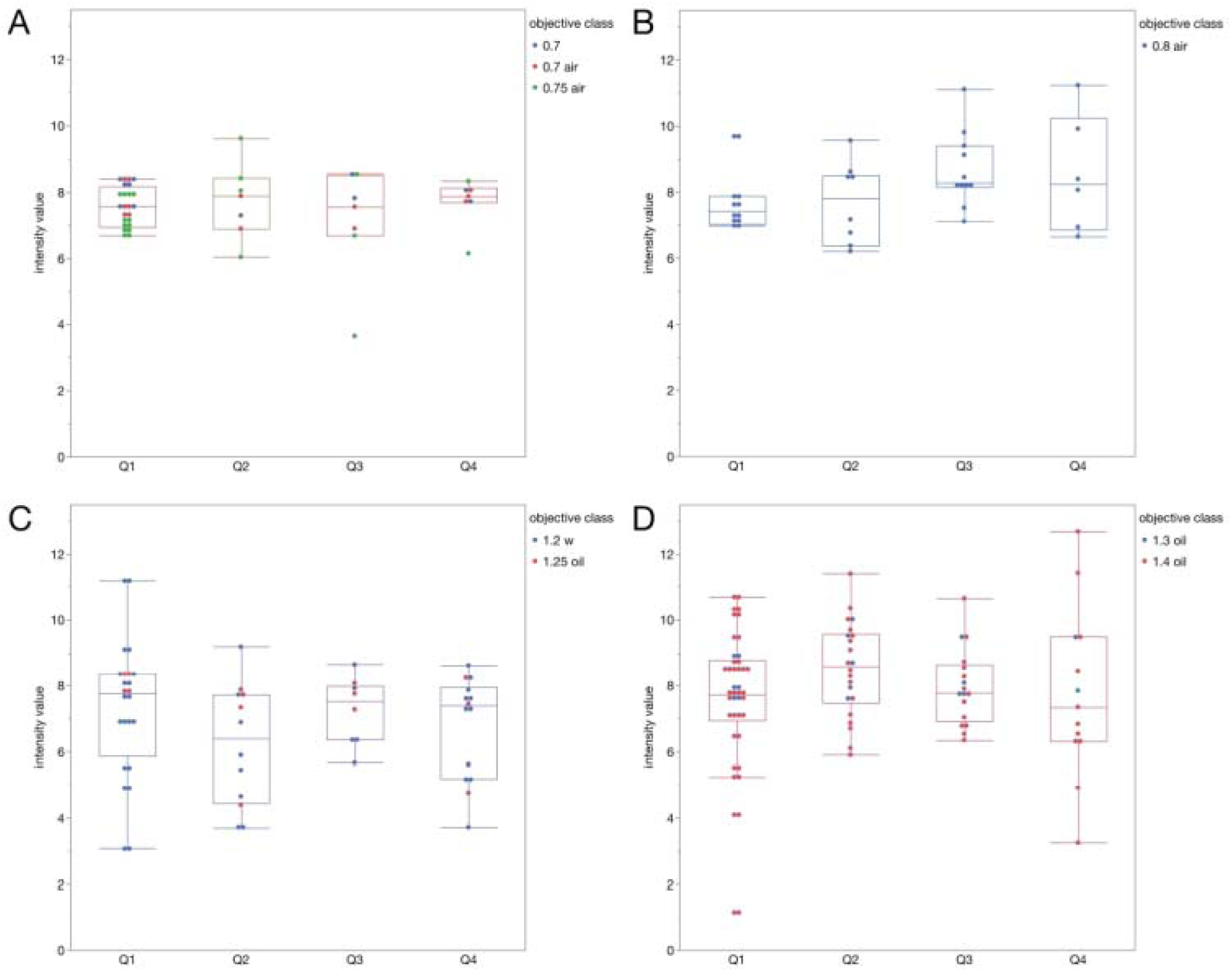
Intensity ratio data broken out by NA. This plot shows the intensity ratio plots for the red microspheres broken out by depth and NA group. 0.7, 0.7 air and 0.75 air (A) 0.8 air (B) 1.2 water, 1.25 oil (C) 1.3 oil, 1.4 oil (D). The ratios are essentially unaffected by depth or NA.

**Supplemental Figure 4.**
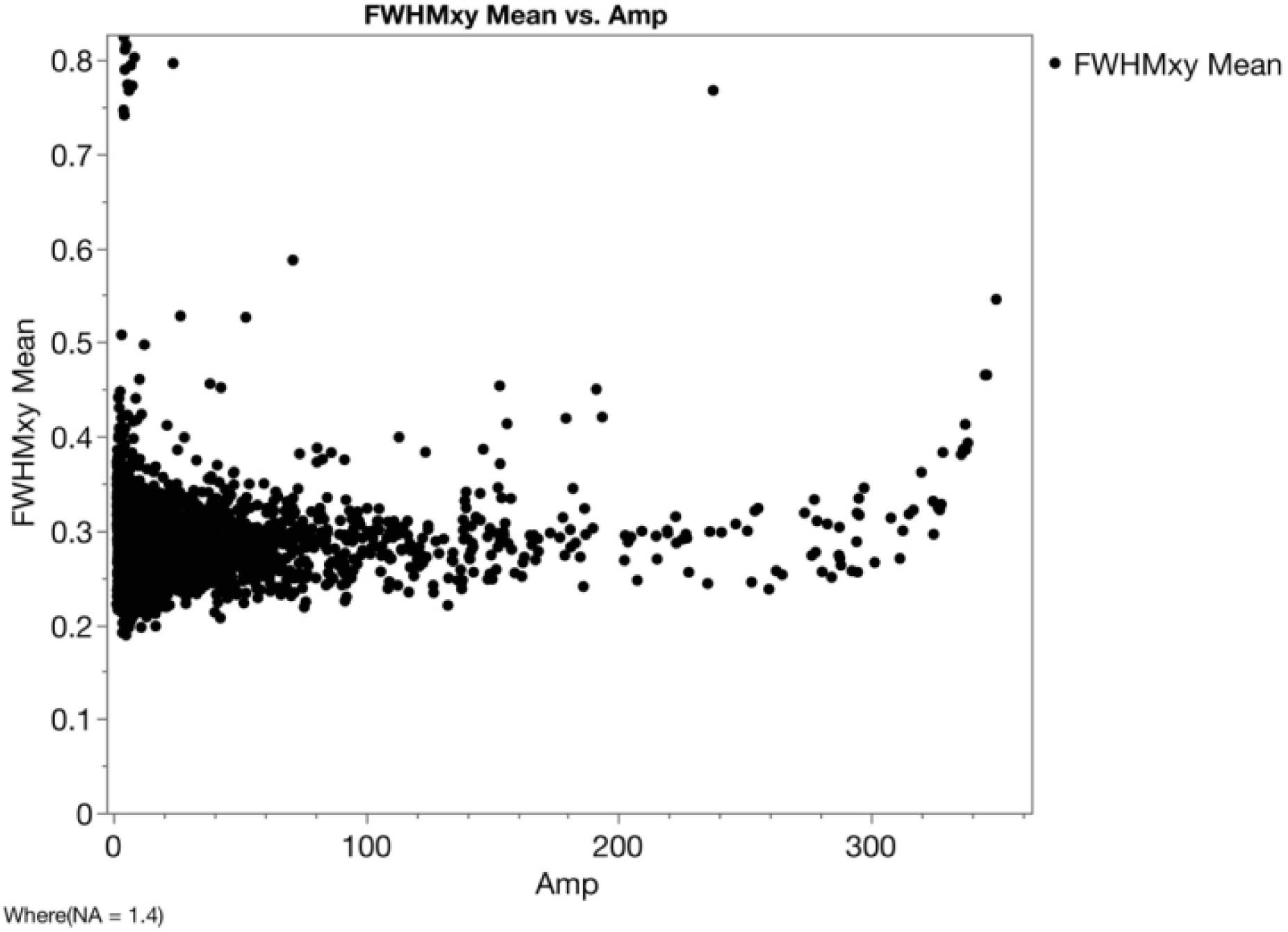
Resolution measurements are robust even for low amplitude microspheres. This plot shows all included FWHMxy-mean values for the 1.4 N.A. objective category (Y-axis) plotted by the signal amplitude (X-axis). This shows that there is not a correlation between the amplitude or intensity of the microsphere and the lateral resolution. This demonstrates that the automated fitting algorithm is robust.

**Supplemental Figure 5.**
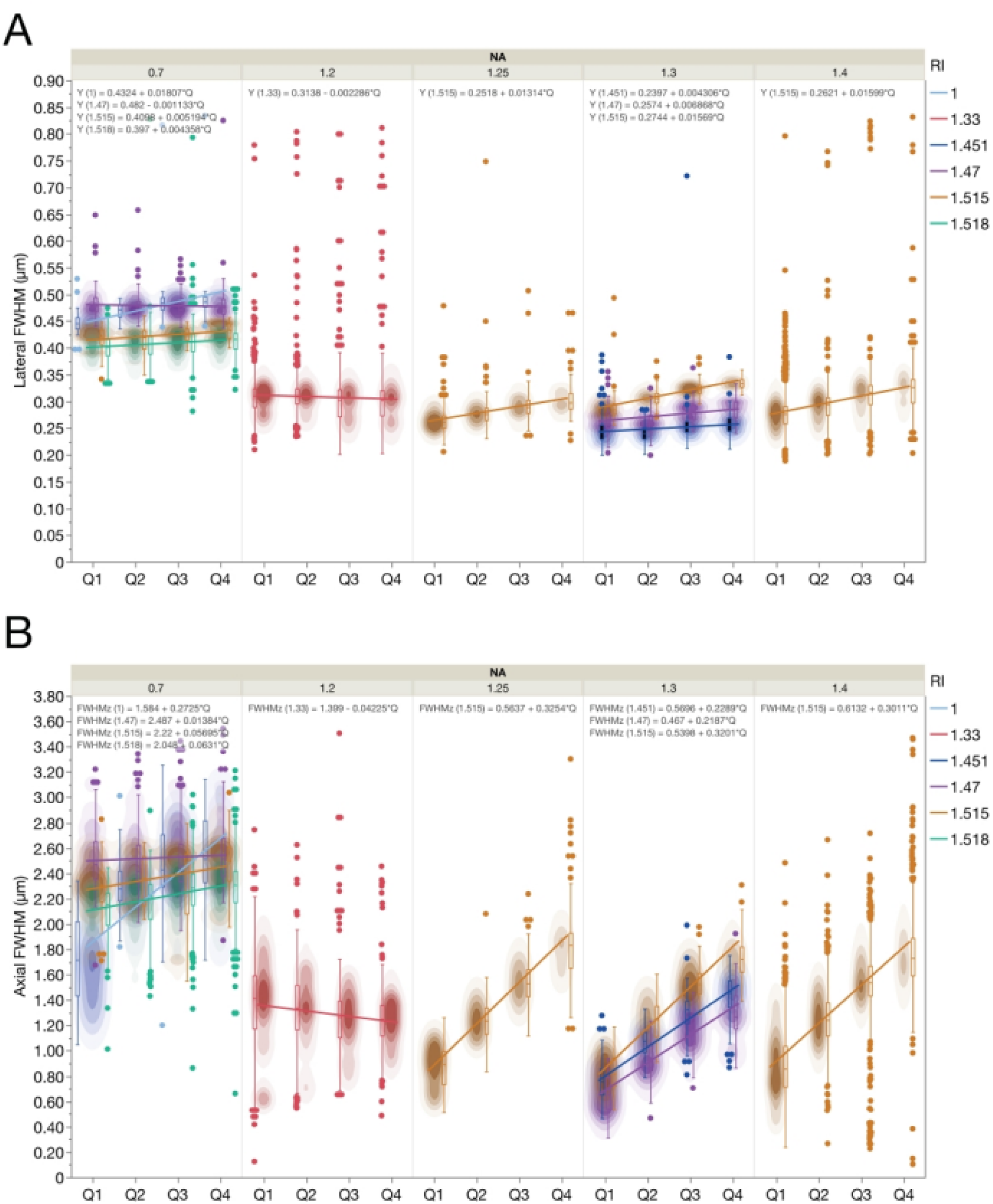
Resolution of microscopes based on PSF curve fitting broken out by refractive index (RI). Lateral (FWHM_xy_) (A) and axial (FWHM_z_) (B) resolution based on curve fitting of PSFs from image stacks of sub-resolution 100 nm diameter green microspheres as a function of depth in the 3D standard CyGel^TM^ sample. This is the same data presented in Figure 2, but now it is broken out by refractive index (RI). The bi- or multi-modal distributions observed in some of the data density (contour plots) for some of the N.A. categories (e.g. N.A. = 1.3 and N.A. = 0.7) can be explained by the refractive index of the lens immersion medium that the lens was designed to be used with relative to the CyGel^TM^. Note that all lens immersion medium/NA categories did not meet the minimum requirements for 100 data points and 3 independent datasets.

